# Braiding Braak and Braak: Staging patterns and model selection in network neurodegeneration

**DOI:** 10.1101/2021.01.21.427609

**Authors:** Prama Putra, Travis B. Thompson, Pavanjit Chaggar, Alain Goriely, for the Alzheimer’s Disease Neuroimaging Initiative

**Affiliations:** Mathematical lnstitute, University of Oxford

## Abstract

A hallmark of Alzheimer’s disease is the aggregation of insoluble amyloid-beta plaques and tau protein neurofibrillary tangles. A key histopathological observation is that tau protein aggregates follow a structured progression pattern through the brain. Mathematical network models of prion-like propagation have the ability to capture such patterns but a number of factors impact the observed staging result, thus introducing questions regarding model selection. Here, we introduce a novel approach, based on braid diagrams, for studying the structured progression of a marker evolving on a network. We apply this approach to a six-stage ‘Braak pattern’of tau proteins, in Alzheimer’s disease, motivated by a recent observation that seed-competent tau precedes tau aggregation. We show that the different modeling choices, from the model parameters to the connectome resolution, play a significant role in the landscape of observable staging patterns. Our approach provides a systematic way to approach model selection for network propagation of neurodegenerative diseases that ensures both reproducibility and optimal parameter fitting.

**Author summary:** Network diffusion models of neurodegenerative diseases are a class of dynamical systems that simulate the evolution of toxic proteins on the connectome. These models predict, from an initial seed, a pattern of invasion called staging. The generalized staging problem seeks to systematically study the effect of various model choices on staging. We introduce methods based on braid diagrams to test the possible staging landscape of a model and how it depends on the choice of connectome, as well as the model parameters. Our primary finding is that connectome construction, the choice of the graph Laplacian, and transport models all have an impact on staging that should be taken into account in any study.

## 1 Introduction

The term ‘neurodegenerative disease’ refers to a family of maladies primarily affecting the brain’s neurons; many neurodegenerative diseases result in cognitive decline and, ultimately, a diagnosis of dementia. A hallmark of such diseases is a large concentration of toxic protein aggregates throughout the brain. These toxic proteins interfere with normal brain functions and are associated with neuronal impairment, neuronal loss, brain atrophy, and overall cognitive decline. Here, we address questions related to model selection for computational studies of Alzheimer’s disease (AD); AD is the most contemporarily prevalent cause of dementia. The practical constraints of clinical AD experiments, most especially in humans, have lead to an interest in the development of mathematical models of pathology evolution; particularly those that can accurately capture large-scale features of AD. Many of these models take advantage of the prion hypothesis which asserts that the progression of proteopathy, in AD, follows from a **prion-like** mechanism.

Prion-like mathematical models can be broadly characterized as either probabilistic [1, 2] or as continuous [3, 4, 5]. Both model types often invoke nonlinearity and are typically discretized, and solved, on either a **simplicial mesh** [6], or on the network defined by the brain’s connectome [3, 4, 5, 7]. Probabilistic and continuous models have their individual advantages. The former can often reveal unique insights in complex datasets while the mathematical structure of the latter can be rigorously analyzed or, in some simple cases, closed form solutions can be derived. Misfolded protein aggregates, in neurodegenerative diseases, often display structured patterns of progression and reproducing these staging patterns is a desirable trait for any mathematical model. Here, we will discuss the impact of model selection, for continuous prion-like mathematical models, with respect to an observed structured staging sequence. As a case study, we will consider the network progression of misfolded tau protein (*τ*P) in AD. We compare results to staging patterns motivated by a recent study 8 that uses a six-stage Braak pattern for *τ*P in AD. However, our braid surface approach is generalizable to more complex hierarchical patterns [9, 10] and well suited to broader investigations; including applications to the study of staging phenomena in other neurodegenerative diseases.

The primary contribution of this manuscript is a methodical, quantitative study of **model selection** features, as they pertain to structured staging, in the setting of a continuous mathematical model of a prion-like proteopathy processes defined on structural networks. This manuscript introduces two novel investigative tools, braid diagrams and braid surfaces, facilitating this process. Our approach is of practical interest for neurodegenerative diseases for two main reasons. First, there is often a lack of the longitudinal radiotracer data, in which staging would be evident, that would provide for accurate, automatic model selection via, for instance, a parameter inference approach. Such studies are important for short-term predictions but they may make model choices based on a limited number of time points and arrive at a model that does not capture a long-term expected staging progression. Second, we show that there are several other factors of a network neurodegeneration model, beyond mathematical parameters, which are not always considered in the model selection process and can play a significant role in staging. In particular, we show that the choice of structural connectome scale, tractography method, thresholding method, **graph Laplacian** weights and, sometimes slight, variation in model parameters can significantly alter the landscape of observed, even accessible, staging patterns.

Our study highlights the nuanced role played by each choice in the network neurodegeneration modeling pipeline; providing important insight, and quantitative tools, that inform model selection.

Even though we choose a particular example of an AD staging as a point of study, the methods and observations are general and can be applied to the study of any hierarchical staging process that can appear in the study of neurodegenerative diseases or other dynamical process on networks. The source code, and connectomes, to create the braid diagrams and braid surfaces discussed in the manuscript are available in the supplementary information.

## 2 Theory and Model

### 2.1 The staging problem

The central problem discussed here is the generalized *staging problem*. For a general dynamical system, we assume that the dynamics on a network are given by

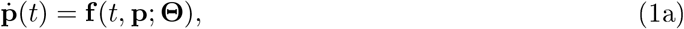

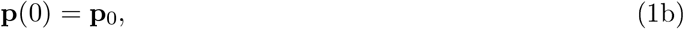

where **p**(*t*) = (*p*_*i*_, … , *p*_*N*_) denotes a normalized and dimensionless quantity, e.g. a normalized concentration 0 ≤ *p*_*i*_(*t*) ≤ 1, evolving on a network *G* = (*V, E*) with nodes *V* and edge set *E*. The quantity *p*_*i*_(*t*) corresponds to the observed concentration in node *υ*_*i*_ and *p*_*i*_(0) is the initial concentration at that node. The quantity **Θ** represents the parameters of the model. In practice, **Θ** represents the parameters of differential equation (1) in addition to other model selection choices such as the edge weighting scheme, the choice of tractography method, which determines graph connectivity and properties that also influence edge weights, or the edge weight thresholding method used to construct the network *G*. We further assume that the dynamics of the system are such that starting from an initial condition, where all concentrations are taken to vanish except one, the system will evolve asymptotically to a state where all concentrations reach their maximal value. It is the spatiotemporal sequence of invasion that we want to characterize.

Let Ω_*j*_, for *j* = 1, 2, … *J* be a non-overlapping collection of nodes; that is, Ω_*j*_ and Ω_*k*_ are disjoint subsets of *V* when *j* ≠ *k*. Let *T* ∈ [0, 1] be an arbitrary, but fixed, threshold value. As **p** evolves, according to (1), the average concentration is computed in each region Ω_*j*_ and the time when Ω_*j*_ first reaches the threshold *T* is recorded. This process produces an ordering of the regions Ω_*j*_ and the ordered sequence 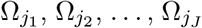 is called an *observed staging pattern*. The *generalized staging problem* is to ascertain the scope of observable staging patterns, subject to variations in **Θ**. The observed staging patterns can then be compared to one or more desirable staging patterns.

### 2.2 A continuous model of *τ*P AD proteopathy on a structural connectome

For our particular application, we model the disease progression on a structural connectome *G* = (*V, E*) with node set *V* given by anatomical regions of interest (ROI) and edge set *E* representing white matter connectivity between these regions. At each node, we define a tau protein concentration, *p*_*i*_, as well as a marker of NFT pathology, *q*_*i*_, and assume the following dynamics:

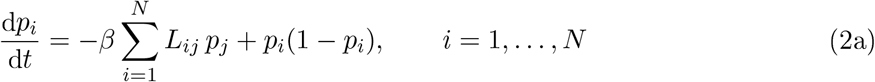

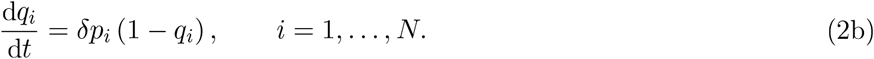

In (2a) the variables *L*_*ij*_ represents the entries of a weighted graph Laplacian **L**, *p*_*i*_ denotes the concentration of seed-competent tau, in node *i*, and *β* a scaled ratio of transmission versus growth (the scaling is chosen so that the coefficient in front of the nonlinear term in (2a) is one). The system (2) is said to be *growth dominated* when *β* ≪ 1 and *diffusion dominated* when *β* ≫ 1. The seed model (2a) is of Fisher-Kolmogorov type and has been considered in several previous studies [4, 11, 7, 6, 12, 13]. In (2b) the variables *q*_*i*_ represent a marker for the local concentration NFT, in a ROI, while *δ* is a coefficient representing an NFT accumulation rate in the presence of *τ*P. This damage model has also been used previously [3, 14, 13]. Since the available staging patterns, observed in clinical imaging data, do not necessarily have a resolution at the level of a single connectome node, we consider *J* regions Ω_1_, … , Ω_*J*_. In each of these regions, we define

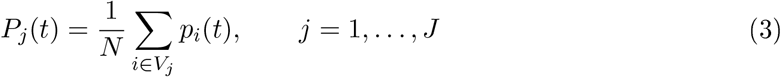

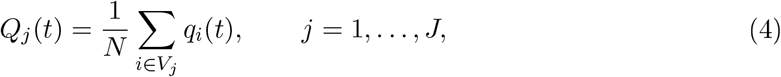

where *V*_*j*_ is the set of nodes defining region Ω_*j*_ with *N*_*j*_ nodes. In all computations, the first region Ω_1_ is defined as the bilateral entorhinal cortex with *N*_1_ nodes and we choose for initial conditions

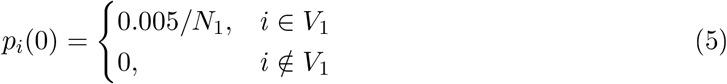

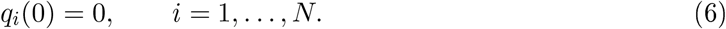

### 2.3 The choice of graph Laplacian

The graph Laplacian used in this work is the *standard graph Laplacian* for an undirected network:

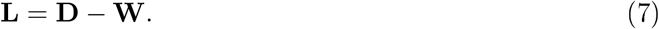

where **W** is the weighted adjacency matrix associated and **D** is the degree matrix, a diagonal matrix with entries

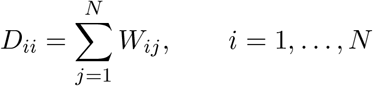

We note that other authors [3, 14, 15, 16, 17] have used normalized forms of the graph Laplacian for their models of neurodegenerative disease progression and a natural question for model selection is the choice of a suitable Laplacian for the underlying process.

A parametrized family of graph Laplacians, encompassing both the standard and normalized forms, is given by

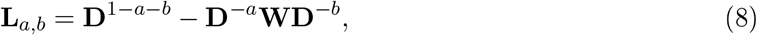

where *a*, *b* ∈ [0, 1] and *a* + *b* ≤ 1, the choice *a* = *b* = 0, yielding the standard graph Laplacian and other popular choices include *a* = −1, *b* = 0, *a* = 0, *b* = 1 and the normali ed graph Laplacian with *a* = *b* = 1/2. The question is to choose, within this family, a graph Laplacian that respects some desirable properties of the transport it is supposed to model. While it is clear that the dynamics, of tau proteins, do not conserve mass overall, since toxic proteins can be either created through aggregation or removed through clearance, the *transport part* of the model should preserve mass. Otherwise, it would require a model assumption to explain how tau proteins are created or removed simply by being transported from one node to the next and how this creation or removal process depend on the degrees of the two nodes and no other mechanism. Since, there is no known such physical mechanism, we are therefore forced to insist on mass conservation by the diffusion part of the model whereas growth and clearance should be modeled by other terms in the model as shown in [4, 18, 5, 19]. We show in the Supplementary Material S1, that the mass-conservation condition is equivalent to the requirement **1** · **L**_*a,b*_ = **0** where **1** = (1, 1, … , 1).

A second condition that we impose on the transport is that transport is driven by difference in concentrations. If the concentration is equal at two nodes, there is no driving force to create an imbalance between these nodes. y analogy with diffusion processes based on *Fick’s law*, we call this condition the *Fick’s condition*. In terms of the graph Laplacian on an undirected network, it reads simply **L**_*a,b*_ · **1** = **0**. We note that there are other possible assumptions when the transport process is viewed as a probabilistic event as shown in [20].

We show in Supplementary Material S1 that the only possible graph Laplacian, in the class defined by **L**_*a,b*_, that satisfies both mass conservation and the Fick’s condition is the standard graph Laplacian **L** = **L**_0,0_. We also show how to generalize this result when regions of different volumes are considered and Fick’s condition suitably generalised.

We want to emphasize that this unavoidable constraint on a physical model of transport does not invalidate previous studies based on the normalized graph Laplacian as these have shown great predictive and explanatory powers when applied to actual patient data. Yet, the choice of using a normalized graph Laplacian is not innocent and should be fully justified as it has some direct implications on the assumptions used in the model, even if these assumptions are not usually given explicitly.

### 2.4 Graph Laplacian weightings

We consider networks *G* = (*V, E*) based on two families of multi-resolution structural connectomes generated from Human Connectome Project (HCP) data (see Methods, Structural connectomes). Each edge, *e*_*ij*_, is associated with two values the number of fibers *n*_*ij*_ constituting the connection between (anatomical region) nodes *i* and *j*; and the fiber length *ℓ*_*ij*_. We consider the choice of weights as part of the model selection and study three possible weights based on the literature: the length-free weighting [3, 15, 14]; the ballistic weighting [4, 18]; and the diffusive weighting [5]. The formulas for these three weighting schemes are listed in Table 1.

**Table 1:** Graph Laplacian weightings for model selection

| Weighting | Formula for $\mathbf{W}_{ij}$ |
| --- | --- |
| Length-free weighting (LW) | $n_{ij}$ |
| Ballistic weighting (BW) | $n_{ij}/\ell_{ij}$ |
| Diffusive weighting (DW) | $n_{ij}/\ell_{ij}^2$ |

### 2.5 Braid diagrams and braid surfaces

A primary contribution of this manuscript is the introduction of braid diagrams and braid surfaces; these powerful tools present a direct and visual assessment of an otherwise complex, nonlinear process evolving in time. We begin by describing a braid diagram. Let *G* = (*V, E*) be a fixed network and suppose that Ω_1_, Ω_2_, *…* , Ω_*J*_ are a fixed set of non-overlapping node regions. Suppose further that *T*_1_, *T*_2_, *…* , *T*_*N*_ are (biomarker) threshold values in the unit interval [0, 1]. A Braid diagram is a graph whose abscissa is the index of the regions Ω_*j*_ and whose ordinate corresponds to the threshold values *T*_*k*_. As a dynamical system, such as (2), evolves on *G*, the time *t*_*j,k*_ at which each region, Ω_*j*_, first achieves each threshold, *T*_*k*_, is recorded. If a given threshold is never achieved in a particular region, the recorded time is prescribed as *t*_*j,k*_ = ∞. The collection of time values, for a fixed threshold index, establishes an ordering of the regions Ω_*j*_. For each threshold, the ordering can then be visualized as a *braid diagram*.

An illustrative example of a braid diagram is shown in Figure 1. Here, the graph consists of four nodes and the regions are simply Ω_*i*_ = {*i*} for *i* = 1, 2, 3, 4. The threshold values are *T*_1_ = 1%, *T*_2_ = 5%, *T*_3_ = 40% and *T*_4_ = 80%. Staging was determined from solving (2a) with ln(*β*) = 3.897 and a synthetic weighting matrix

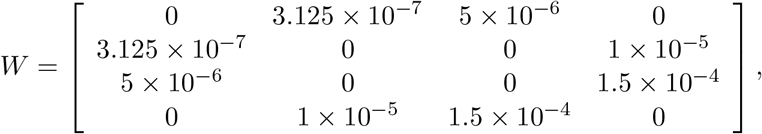

which corresponds to the DW scheme (Table 1) for illustrative edge lengths *ℓ*_1,2_ = *ℓ*_2,1_ = 40 and all other edge lengths equal to 20. This simple example shows, for instance, that the regions achieve the threshold *T* = 5% in the order Ω_1_ → Ω_3_ → Ω_2_ → Ω_4_. However, for the threshold *T* = 40%, the observed ordering of regions changes to Ω_1_ → Ω_3_ → Ω_4_ → Ω_2_. These two observed staging patterns can be expressed by the abbreviated notation I→III→II→IV and I→III→IV→II, respectively. A braid diagram is useful for considering observed staging for a fixed set of model parameters; for instance, for a fixed value of *β* in (2).

**Figure 1:**
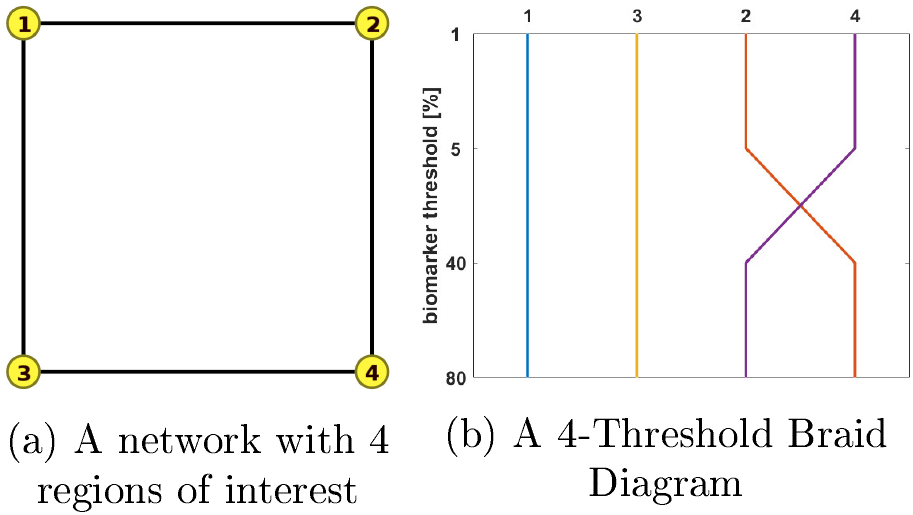
A network with 4 regions of interest (left). A braid diagram (right) generated from an illustrative configuration of fibers (*n*_*ij*_), edge lengths (*ℓ*_*ij*_), DW scheme and dynamics (*β*).

We are also interested in staging outcomes as the system parameters are varied. This information is contained in a *braid surface* that generalizes a braid diagram. For each possible staging, we assign a color. The braid surface is a two-dimensional plot that assign for each value of one parameter and one biomarker the corresponding staging color. Computationally, this surface is generated by a simple algorithm. First, one discretizes the continuous values of a parameter, such as *β* in (2), of interest. Then, for each discretized parameter, the underlying system (e.g. (2)) is solved, a braid diagram is constructed and the observed staging pattern is determined for each threshold value; if an observed staging pattern has not yet been encountered, it is added to a list. At the end of the process, every pairing of discrete parameter and threshold has been assigned to an observed staging pattern. The set of observed staging patterns are assigned to colors and these colors are plotted to visualize the braid surface. The *τ*P seed staging braid surface for the example network, and weighting matrix, of Figure 1 is shown in Figure 2. The green 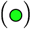 corresponds to the *β*-parameter, and threshold, region where a I→III→II→IV staging pattern is observed while the red 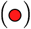 corresponds to the parameter region where a I→III→IV→II is produced. The braid surface shows that the staging dynamics, for the four-vertex system with diffusive weights, are simple; we will observe complex staging dynamics on structural connectome graphs of the brain.

**Figure 2:**
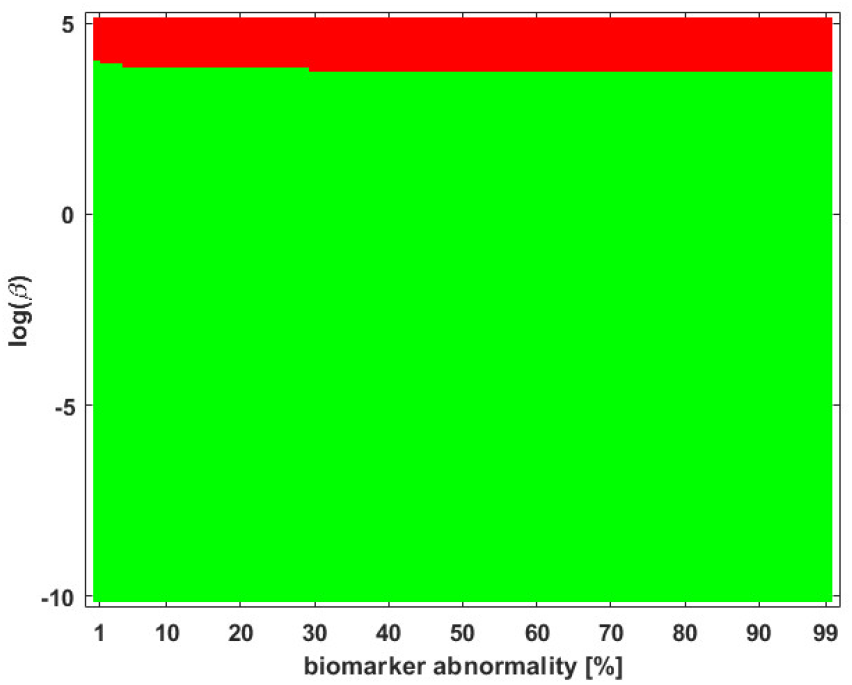
A braid surface showing the *τ*P seed staging dynamics for the simple network of Figure 1 as *β* varies. Two distinct staging patterns are produced for the (diffusive) weights considered.

### 2.6 Hierarchical staging of *τ*P in AD

The problem of the structured staging of *τ*P NFT in AD has been well studied. The most famous study being the seminal work of Braak and Braak [21, 22] that looked at the progression through six regions. The six-region progression view is still in contemporary use [1, 8] though some authors have also proposed refinements to ten regions [9]. More recently, the hierarchical progression of flortaucipir tracer has also been studied for progression patterns; authors have studied SUVR progression using a canonical six-region [23] and an extensive 25-region [10] pattern.

Much of the staging literature refers to the evolution of *τ*P NFT or of SUVR quantities; though a recent study [8] has advanced the notion that *τ*P seeds precede NFT pathology in a similar structured manner. The model that we study, i.e. (2), tracks the progression of both *τ*P seeds (via (2a)) and *τ*P NFT (via (2b)). Therefore, we adopt the staging regions given by [8]; we will observe how the many, sometimes surprising, aspects of model selection can strongly influence the set of observed regional progressions for a computational model of proteopathic *τ*P staging in AD. The same method can easily be generalized to the study of other staging regions such as those advanced in [1, 2, 10] , or [9].

## 3 Results

We studied the staging problem for the model (2) of *τ*P proteopathy in AD, using the braid surface approach. To study this problem, a five-region adaptation, to match the structural connectome ROI, of the six regions used in [8] was adopted and is shown in Table 2.

**Table 2:** Connectome regions used in the staging problem.

| Stage | Connectome ROI | Stage | Connectome ROI |
| --- | --- | --- | --- |
| Region I | entorhinal cortex | Region II | hippocampus |
| Region III | parahippocampal gyrus | Region IV | rostral anterior cingulate, caudal anterior cingulate |
| Region V | cuneus, pericalcarine cortex, lateral occipital cortex, lingual gyrus |  |  |

We refer to staging progression, determined by a concentration threshold 0% < *T* ≤ 100%, through these five regions with Roman numerals and use the I→II notation. To present the braid surface results, we first enumerate all observed regional staging patterns and assign each one a color as shown in Table 3.

**Table 3:**
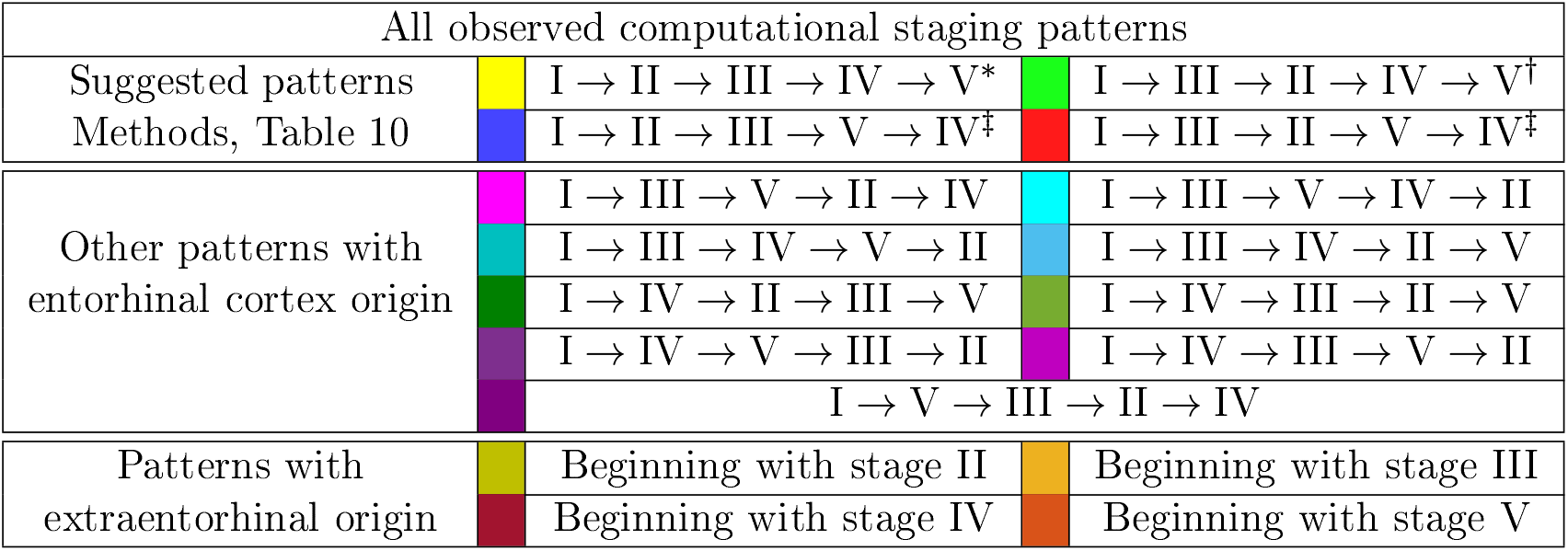
Color coding for braid surfaces. * Progressive computational Braak staging. ^†^ Computational staging suggested by SUVR data. ^‡^ Additional potential computational staging, *τ*P seeding.

| All observed computational staging patterns |  |  |  |  |
| --- | --- | --- | --- | --- |
| Suggested patterns | | $I \rightarrow II \rightarrow III \rightarrow IV \rightarrow V^*$ | | $I \rightarrow III \rightarrow II \rightarrow IV \rightarrow V^\dagger$ |
| Methods, Table 10 | | $I \rightarrow II \rightarrow III \rightarrow V \rightarrow IV^\ddagger$ | | $I \rightarrow III \rightarrow II \rightarrow V \rightarrow IV^\ddagger$ |
| Other patterns with entorhinal cortex origin | | $I \rightarrow III \rightarrow V \rightarrow II \rightarrow IV$ | | $I \rightarrow III \rightarrow V \rightarrow IV \rightarrow II$ |
| | | $I \rightarrow III \rightarrow IV \rightarrow V \rightarrow II$ | | $I \rightarrow III \rightarrow IV \rightarrow II \rightarrow V$ |
| | | $I \rightarrow IV \rightarrow II \rightarrow III \rightarrow V$ | | $I \rightarrow IV \rightarrow III \rightarrow II \rightarrow V$ |
| | | $I \rightarrow IV \rightarrow V \rightarrow III \rightarrow II$ | | $I \rightarrow IV \rightarrow III \rightarrow V \rightarrow II$ |
| | | $I \rightarrow V \rightarrow III \rightarrow II \rightarrow IV$ | | |
| Patterns with extraentorhinal origin |  | Beginning with stage II |  | Beginning with stage III |
|  |  | Beginning with stage IV |  | Beginning with stage V |

From the original point of view [21] of *τ*P Braak staging, one would expect that the I → II → III → IV → V progression is a likely candidate of interest; at least for NFT. However, based on ADNI data, we also identified three other observed progression sequences that may be of interest for both *τ*P seeds and NFT. The set of identified staging patterns of potential interest are called Suggested Patterns and summarized in Table 3 (top).

### 3.1 Connectome preparation significantly alters observed staging patterns

We compared two different types of connectome tractography methods; both *deterministic tractography*, prepared by [24] using the MRtrix software, and *probabilistic tractography*, prepared for this study using the FSL software. These generating methods used parcellation ROIs defined by the Lausanne multi-resolution atlas and constructed from HCP data. The braid surfaces for the deterministic connectomes and the probabilistic connectomes are, in general, quite different. However, there are some notable similarities. For instance, both connectome types show great sensitivity as the parameter *β* increases to large values. In this diffusion-dominated regime, the observed staging pattern becomes non-distinct with multiple overlap for small changes of parameters. Another common feature is that there are large areas, of *T* and *β* parameter space, where the *τ*P seed staging and *τ*P NFT staging are consistent; meaning that both display the same staging pattern. A final notable similarity is that both connectome types are able to produce the progressive Braak 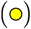 pattern for NFT staging.

Despite these similarities, there are also clear differences between the staging behavior observed on the deterministic and probabilistic connectomes. In particular, we examined ADNI SUVR data subject to an ROI selection from a recently published study [8] on *τ*P seeding along the Braak pathway. The SUVR staging pattern suggested 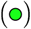 by our analysis appears prominently on the deterministic connectomes (Figures 5 and 4) but is nearly absent on the probabilistic connectome counterparts (Figures 5 through 9). In fact, the deterministic connectomes can express all four suggested staging patterns (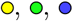 and 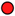) at high resolution for both *τ*P seeds (Figure 3) and NFT (Figure 4) while the probabilistic connectomes almost always express fewer patterns.

**Figure 3:**
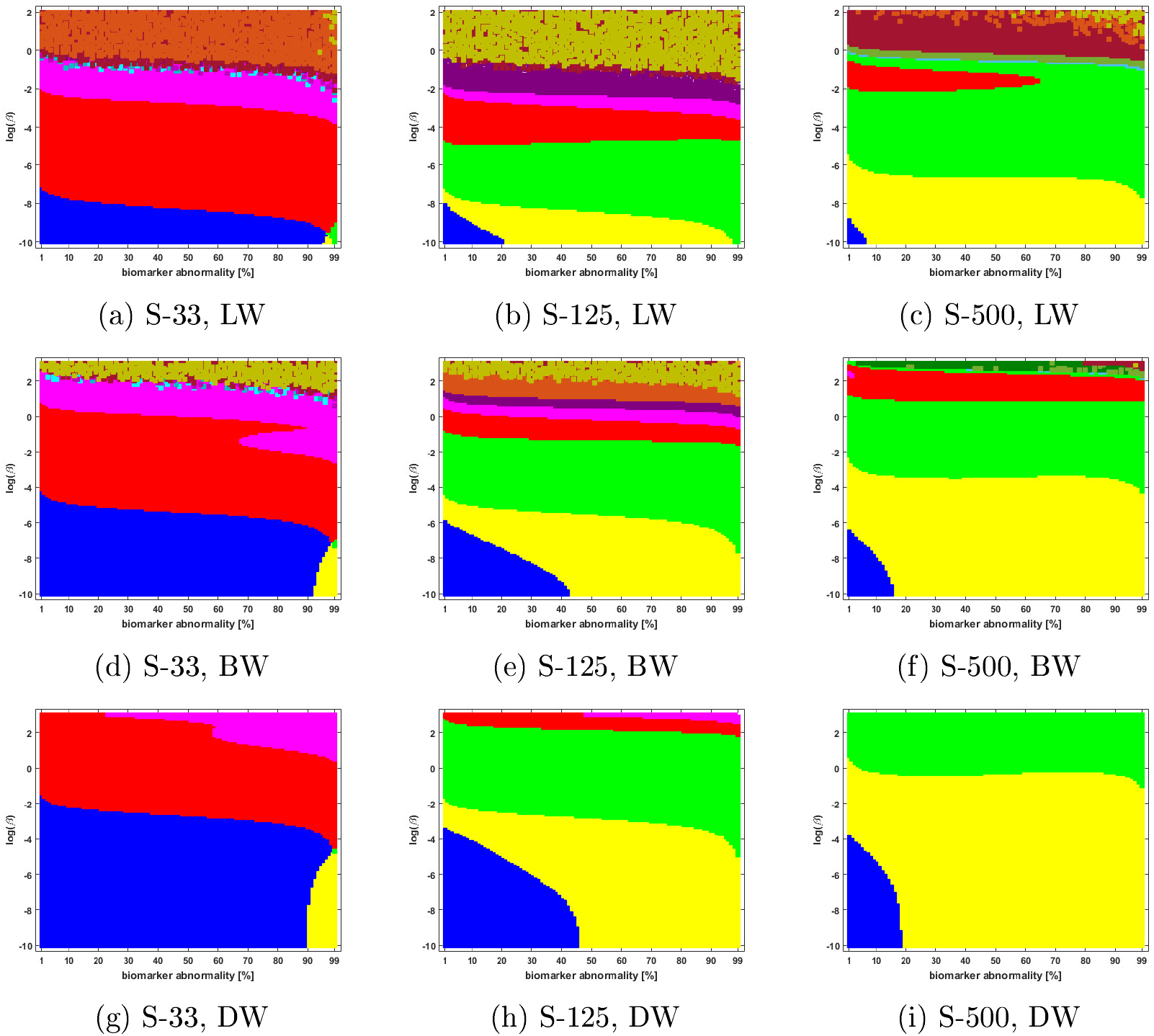
Observed computational (deterministic) connectome *τ*P seed staging; length-free (top), ballistic (middle) and diffusive (bottom) weighting schemes. The *x*-axis determines the biomarker abnormality threshold 1% < *T* ≤ 100% and the *y*-axis corresponds to −10 ≤ ln(*β*) ≤ 2 for the parameter *β* in (2a).

**Figure 4:**
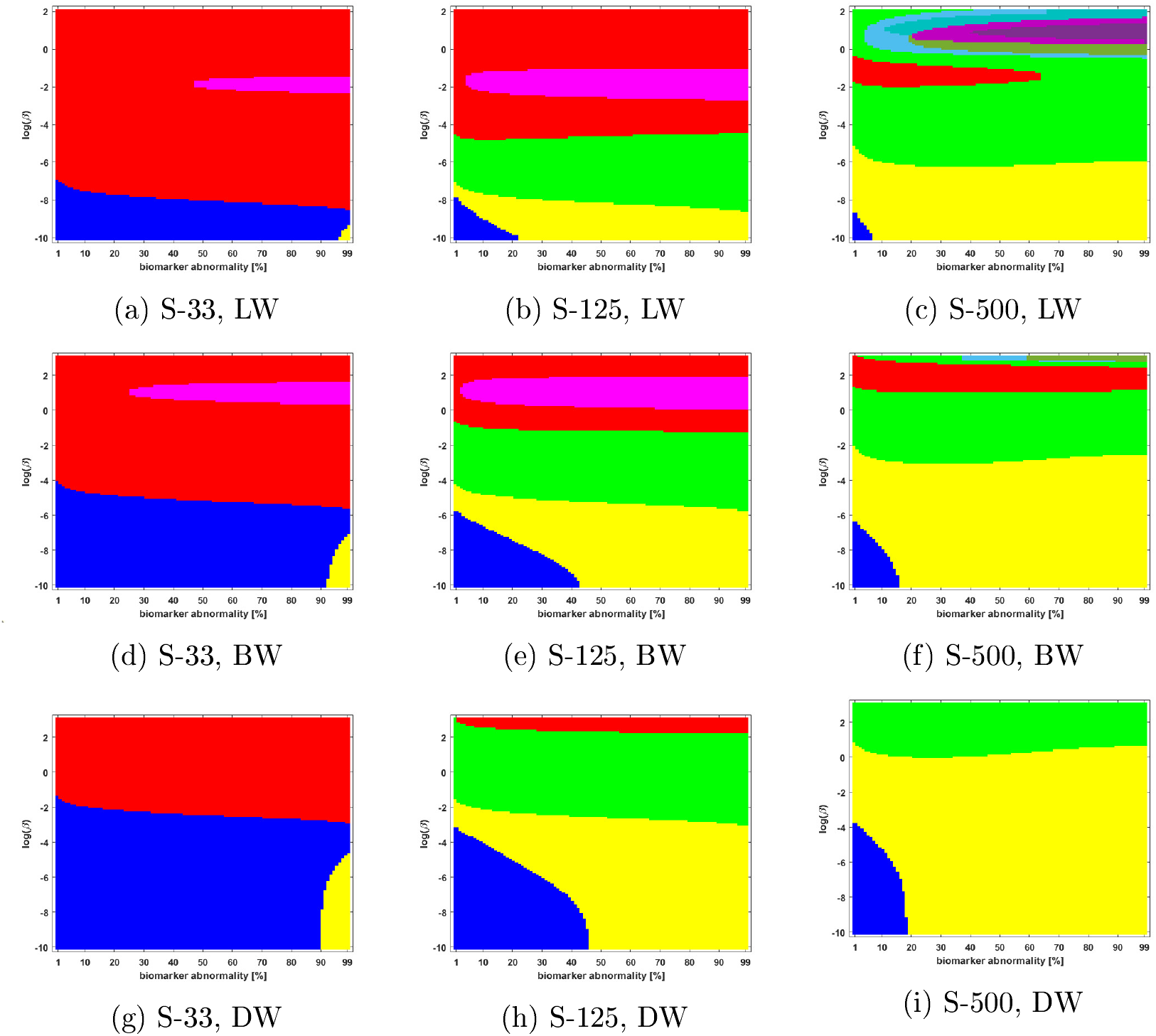
Observed computational (deterministic) connectome *τ*P NFT staging with *δ* = 1 in (2b). Figure order, axis labels and axis ranges are identical to those of Figure 3.

**Figure 5:**
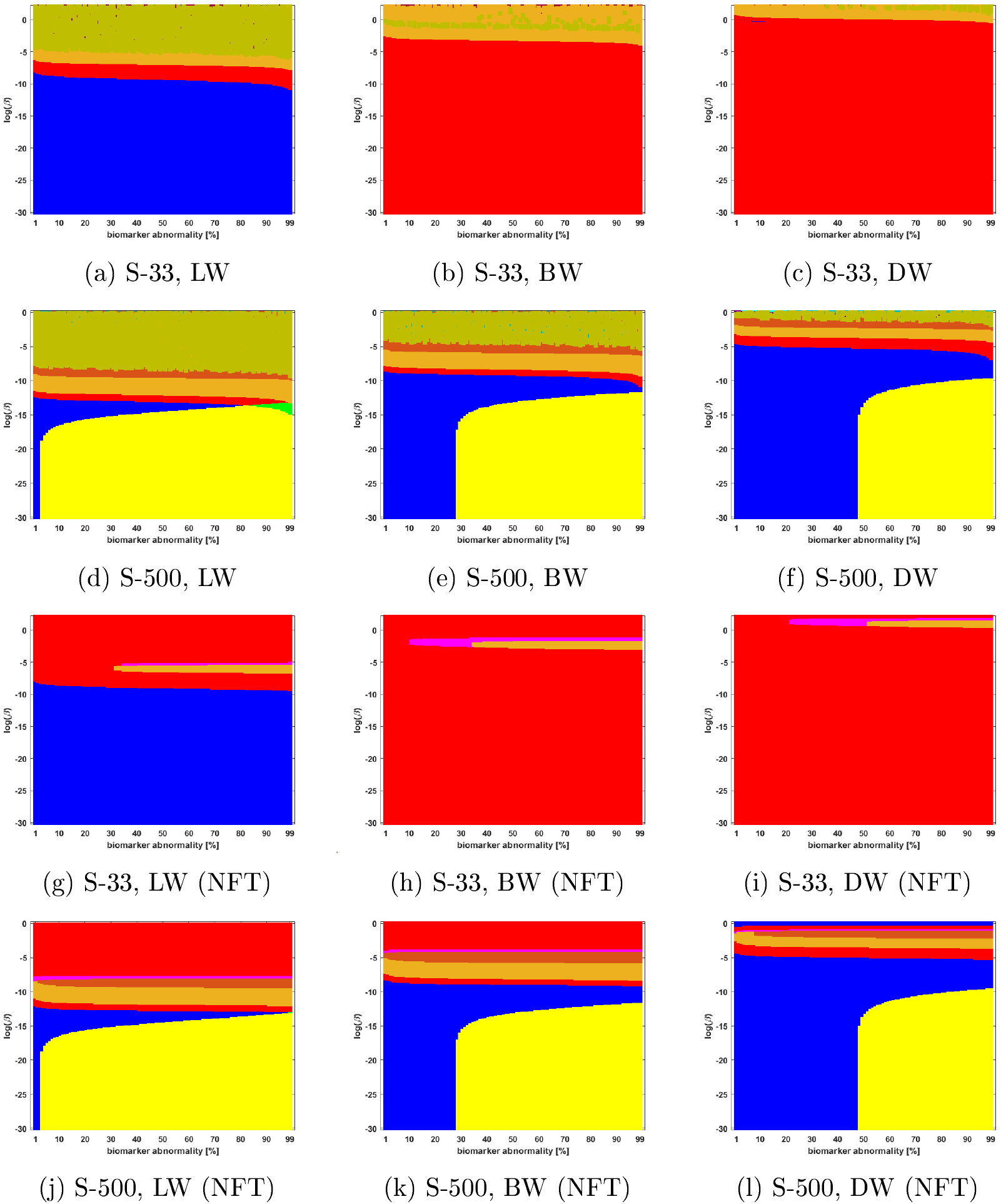
Observed computational (probabilistic) connectome *τ*P seed staging (top two rows) and *τ*P NFT staging (*δ*=1, bottom two rows). Density filter (DF) thresholding at a threshold of 2 × 10^−1^. The *x*-axis determines the biomarker abnormality threshold 1% ≤ *T* ≤ 100% and the *y*-axis corresponds to −30 ≤ ln(*β*) ≤ 0 for the parameter *β* in (2a).

We also observed (Figures 5 through 9 and supplementary information S2) significant qualitative differences in the braid surface landscapes when various thresholding methods were used in preparing the probabilistic connectomes (Table 4). Therefore, we conclude that connectome preparation plays an important role in all downstream results and is an integral part of model selection.

**Table 4:**
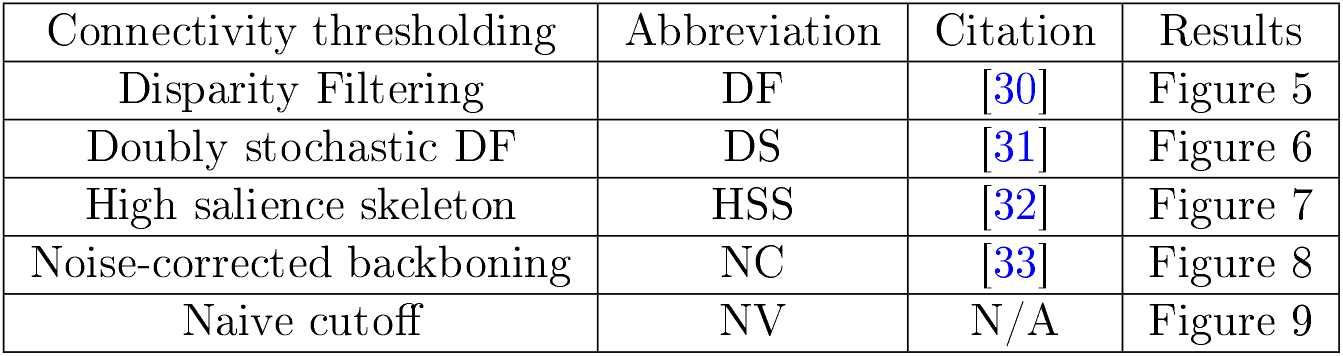
Thresholding methods for probabilistic connectomes

### 3.2 The graph Laplacian weighting scheme modifies observed staging paradigms

Several graph Laplacian weighting schemes have been suggested to model disease dynamics. But, it remains unclear as to which of these weighting choices is most biologically sound. Our results suggest that this choice also affects the observed computational staging patterns in two ways. The first of these concerns *τ*P seed staging within the parameter regime where diffusion takes a more prominent role (i.e. large values of *β*). The move from length-free (LW) to ballistic (BW) weights tends to decrease the surface area where the observed computational staging is sensitive to small changes the in biomarker abnormality threshold; this surface area is again reduced when moving from ballistic (BW) to diffusive (DW) weights. This can be seen in both the deterministic (Figure 3) and probabilistic (Figures 5 through 9, top two rows) braid surface results. In this sense, the length dependence in the weights ‘stabilizes’ the *τ*P seed staging, appearing with an increase in *β*, in (2a).

The second general observation concerns the prevalence of the progressive Braak 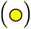 and SUVR - suggested 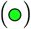 staging patterns. In the former case, the general trend is that the choice of a DW graph Laplacian tends to increase the prevalence of the progressive Braak 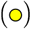 NFT staging (Figure 4 and Figures 6 through 9, bottom two rows). However, we remark that this observation can depend on the choice of thresholding technique as we observed in the case of the density filtered probabilistic connectome where the LW had this effect (Figure 5, bottom two rows and supplementary information S2 Figure 3, bottom two rows). However, if one were comparing computational results directly to SUVR staging (e.g. [1, 2]), using the ROI of [8], the observation is more complex. In the case of deterministic streamlining, the prevalence of the observed SUVR-suggested staging 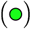 can decrease, within the ranges of *β* we considered, as one moves from LW to BW and again to DW (Figure 4, right-most column). However, we do note that a number of undesirable staging patterns are observed, for LW, when ln(*β*) > 0 on the highest resolution connectome (Figure 4c); thus, extra consideration should be given to the mathematical model parameters in this case. For probabilistic connectomes, the appearance of the SUVR-suggested staging pattern 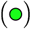 depends more, in general, on thresholding technique and the parcellation resolution; the choice of weights had only a weak impact in this specific regard. Our results suggest that, in the absence of a clear biological impetus for selecting particular graph Laplacian weights, a braid surface analysis can play an important, practical role in selecting amenable weights for a particular study.

**Figure 6:**
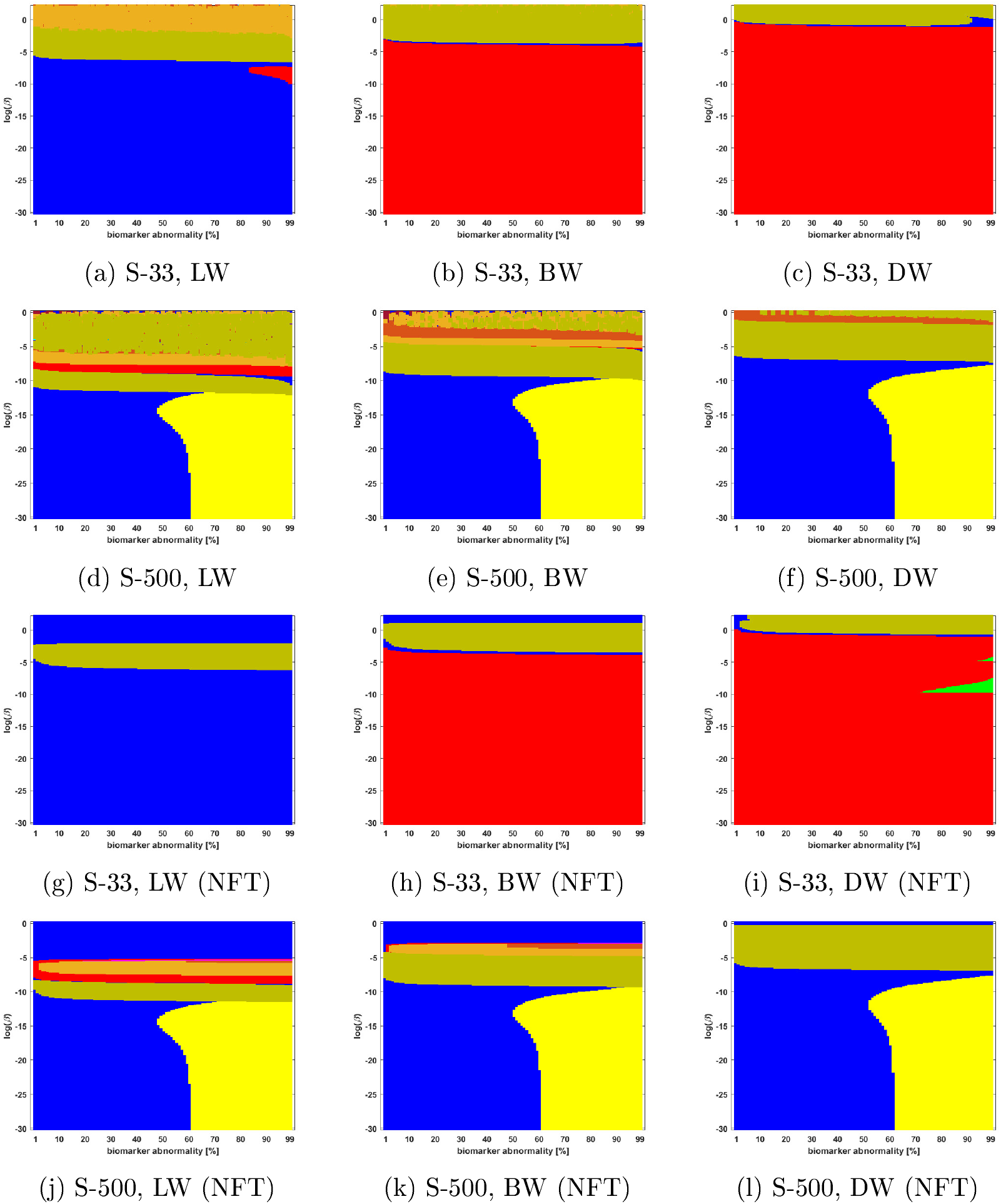
Observed computational (probabilistic) connectome *τ*P seed staging (top two rows) and *τ*P NFT staging (*δ* = 1, bottom two rows). Doubly stochastic thresholding at a threshold of 1 × 10^−2^. Axes coincide with those of Figure 5

**Figure 7:**
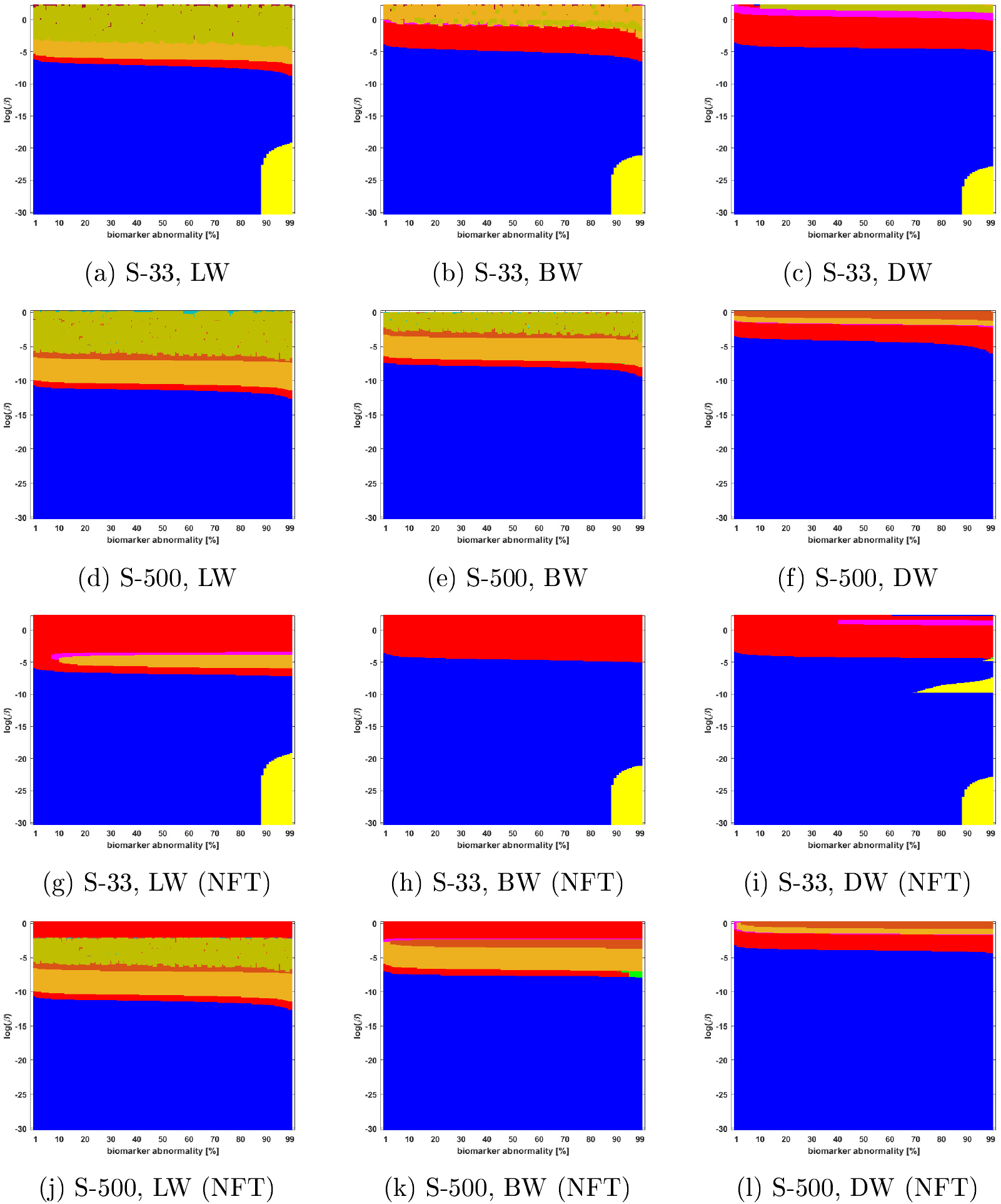
Observed computational (probabilistic) connectome *τ*P seed staging (top two rows) and *τ*P NFT staging (*δ* = 1, bottom two rows). High salience skeleton at a threshold of 1 × 10^−1^. Axes coincide with those of Figure 5

**Figure 8:**
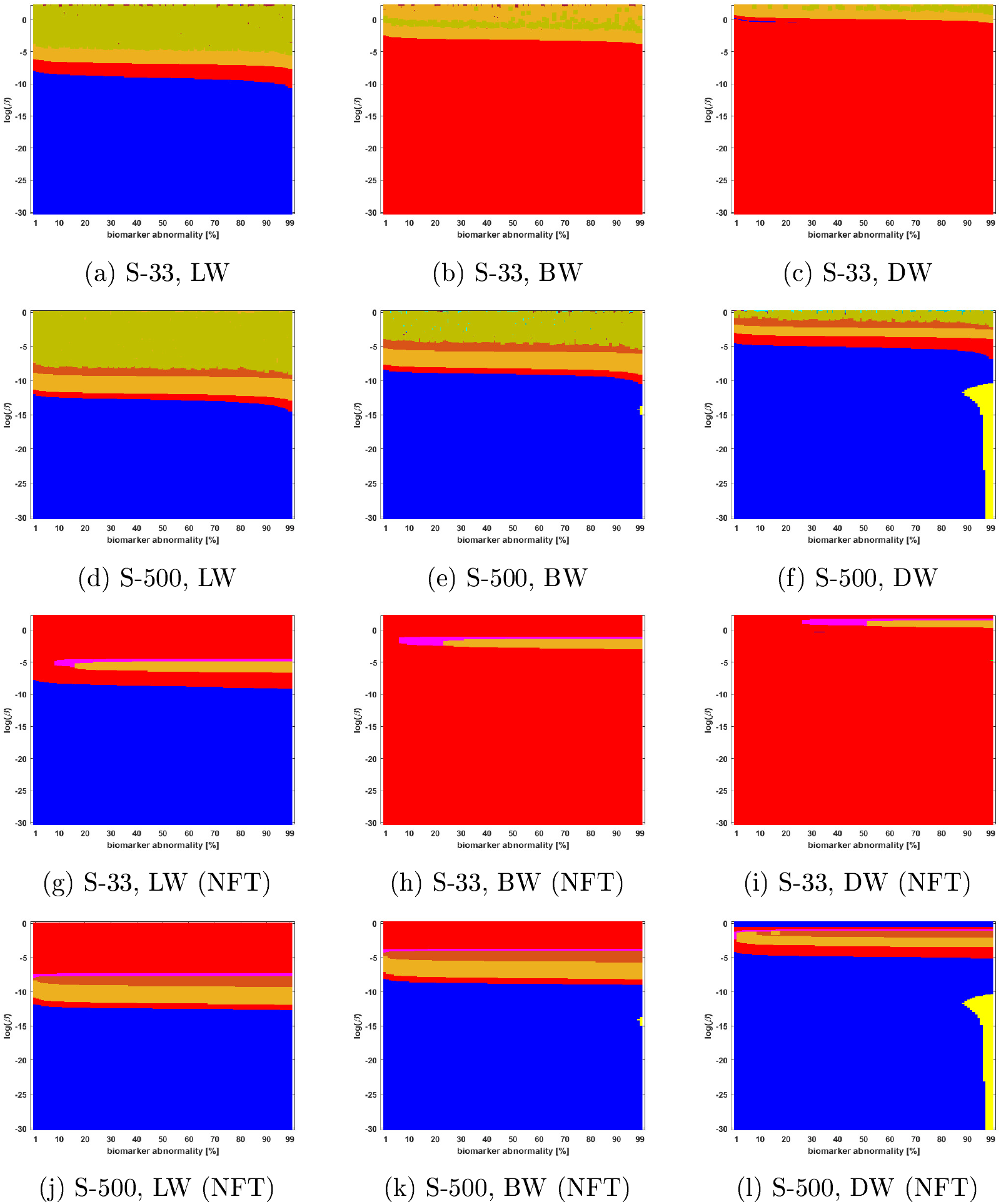
Observed computational (probabilistic) connectome *τ*P seed staging (top two rows) and *τ*P NFT staging (*δ* = 1, bottom two rows). Noise corrected backbone at a threshold of 2.32. Axes coincide with those of Figure 5

**Figure 9:**
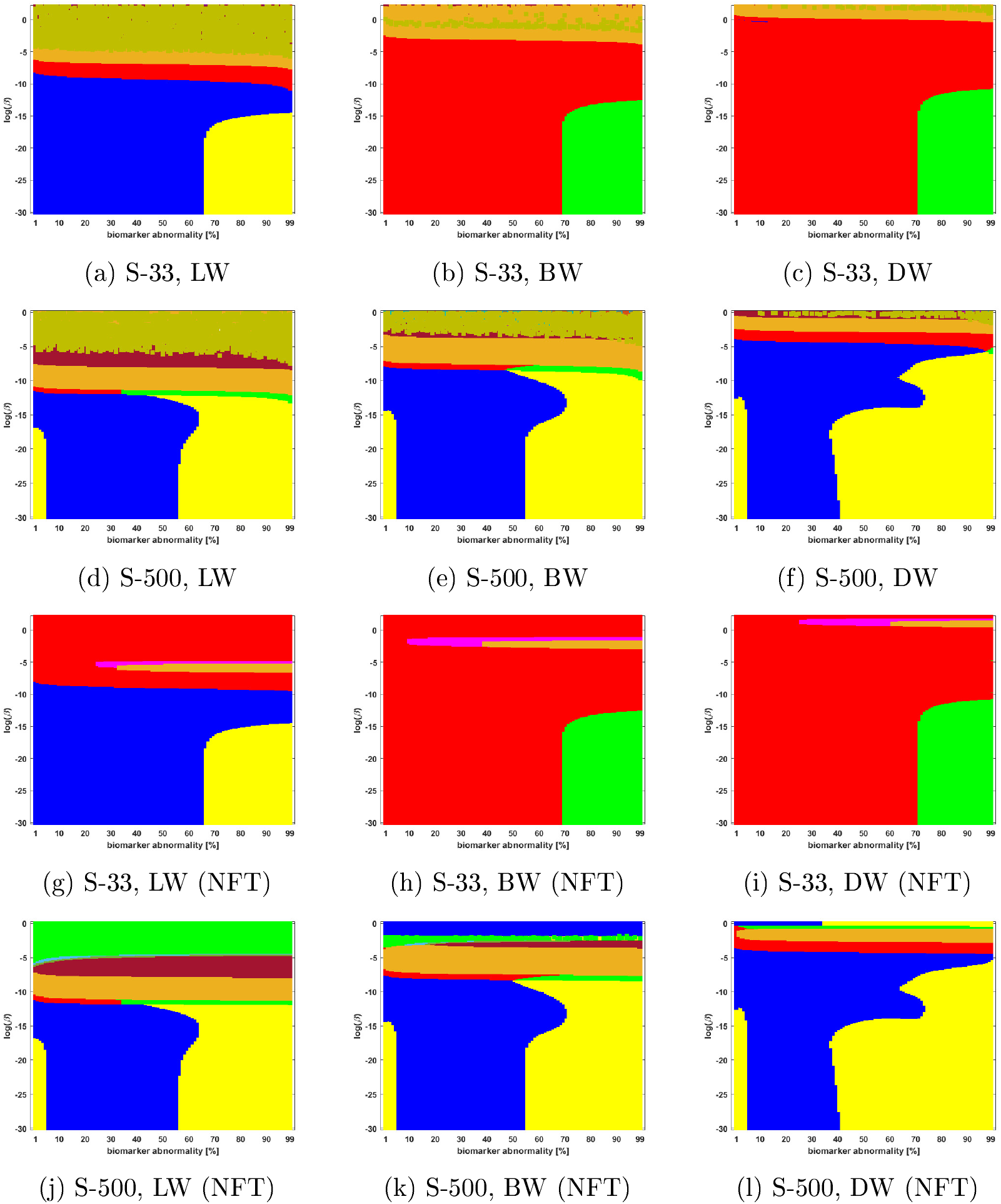
Observed computational (probabilistic) connectome *τ*P seed staging (top two rows) and *τ*P NFT staging (*δ* = 1, bottom two rows). Naive thresholding at a threshold of 1 × 10^−2^. Axes coincide with those of Figure 5

### 3.3 Connectome resolution has variable effects on observed staging

Network neurodegeneration studies have evolved systems, such as (2), on both low-resolution [3, 14, 4, 13, 7] and high-resolution [5] connectomes. We studied how connectome resolution alters observed computational staging patterns. In the case of connectomes generated with deterministic streamlining, increasing the connectome resolution improved the overall observed staging results. For instance, we see a stabilizing effect on *τ*P seed staging, when ln(*β*) > 0, (columns of Figure 3) with increased connectome resolution. Moreover, the combined area of the progressive and SUVR - suggested staging patterns of interest (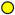 and 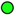, respectively) tend to increase in prominence with increased resolution; conversely, the transposed prefix stagings (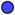 and 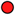) are more pronounced at lower resolutions. This observation also holds for NFT staging (Figure 4).

In the case of probabilistic streamlined connectomes, the thresholding method and the effect of resolution are intertwined (Figures 5 through 9). It is invariably clear that resolution has a pronounced effect on observed staging but those effects varied depending on the thresholding method. In general, the low resolution connectomes consisted primarily of a single staging pattern (typically either 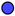 or 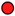) when growth was sufficiently strong (circa ln(*β*) < −5) and increasing the connectome resolution increased heterogeneity in addition to introducing the progressive Braak staging pattern 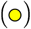.

Together, our results show that connectome resolution plays a deciding role in the landscape of observable staging patterns. In particular, the seemingly ubiquitous practice of the use of low-resolution connectomes may partially explain discrepancies between modeling results and comparisons to imaging data in the latter Braak regions (e.g. IV and V).

### 3.4 Braid surfaces for deterministic connectomes

Deterministic connectomes were constructed using deterministic streamlining (see Methods). We report the observed staging for both *τ*P seeds (Figure 3) and for NFT (Figure 4). We present these results for the lowest (Scale-33), median (Scale-125) and highest (Scale-500) resolution connectomes in addition to all three graph Laplacian weighting schemes. Both figures summarize results for the model parameter value *β* in the range −10 ≤ ln(*β*) ≤ 2. Further results are reported in the supplementary material (S2).

### 3.5 Braid surfaces for probabilistic connectomes

Probabilistic connectomes were constructed from HCP data. We focus on the effects of the parameter *β* and of several connectivity thresholding methods on observed staging results. For brevity, we present only one thresholding level per thresholding method but other thresholding levels are presented in the supplementary information (S2).

## 4 Discussion

In the context of studying prion-like spreading of misfolded proteins in AD, different studies have made use of different model choices. For instance, within the class of reaction-diffusion models such as (2), several types of graph Laplacian weights have been used; including those that are free of the influence of length [3, 14, 25] weights that model prion-like transport as a velocity [4, 18] or as diffusive [5] in nature. Various connectome resolutions, for the Lausanne parcellation, have also been used; both low resolution [3, 14, 25, 4, 18, 7, 13] and high resolution [5]. Several works have compared, or fitted, spreading models to atrophy [14] and SUVR data [25, 1, 2, 7]. However, to our knowledge, a systematic methodical investigation into the general staging problem for AD has not been advanced, even for the simpler class of prion-like progression models such as (2).

In this manuscript we have undertaken a methodical investigation of the generalized staging problem for a simple network model of AD; this important problem concerns the ordered progression of a marker of interest propagating over a network. Using an adaptation of a six-stage sequence of ROIs [8] along the Braak pathway, we have systematically investigated how various aspects of model selection, including those mentioned above, may alter the progression of *τ*P seeds and NFT across the connectome. To do so, we have introduced, and applied, the novel tools of braid diagrams and braid surfaces. Though we have focused on a continuous network neurodegeneration model, our method also applies to probabilistic spreading models as well [26, 1].

### 4.1 Connectome construction limits realizable staging patterns

Our findings have several implications for model selection. The most surprising is that the particulars of the connectome itself play an important role in the staging problem. That is, when studying *τ*P progression, the construction of the structural connectome should be considered as an inextricable part of the model. To illustrate this, consider two particular staging patterns; the progressive Braak pathway NFT staging pattern 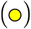 and the staging pattern suggested 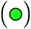 by our ADNI flortaucipir data analysis. It is worth noting that the difference between the progressive Braak 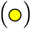 versus the SUVR-suggested 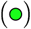 staging is whether a biomarker first appears in the hippocampus (II) or the posterior parahippocampal gyrus (III). In [8], the histopathologically identified Braak stage II brains expressed *τ*P seeding in the parahippocampal gyrus (III) at a slightly higher average level than in the hippocampus; potentially explaining the difference between the SUVR suggested 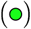 and progressive 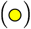 staging patterns. That is, 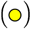 and 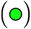 may essentially reflect the same effective staging but from two different points of view. Only the 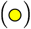 pattern was reliably expressed on the often-used, low-resolution (Scale-33) structural connectome constructed with deterministic streamlining. However, both patterns appeared in the high resolution (Scale-500) case (Figures 3 and 4, left column v.s. right column). Conversely, the high resolution connectomes generated with probabilistic tractography, and subsequently thresholded for sparsity, almost never produced the 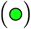 pattern suggested by our ADNI data analysis (Figures 5 through 9, second and fourth rows); though they did produce the progressive staging 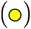. The SUVR pattern 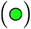 did, however, appear robustly for the Scale-33 connectome when naive thresholding was used (Figure 4). For the Scale-500 naively thresholded connectome, the 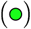 pattern does appear but is only consistent with seed staging in a very small region of parameter and threshold space (see Figure 4, second and last rows). Figure 4 suggests that naive thresholding may be a promising strategy for probabilistic connectomes; however, we found that the 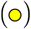 and 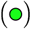 patterns were sensitive to the threshold used (see Supplementary Material S2) and were almost completely absent, at both scales, for the nearby thresholds we examined.

The consequences of these observations are non-trivial. For instance, a study comparing the NFT model (2b) to ADNI flortaucipir data on the low resolution probabilistic tractography connectomes, thresholded with a density filter, would not be capable of achieving the staging pattern 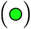 obtained by our SUVR analysis (see Figure 5, third row); invariably leading to a misfit between model and data. Overall, our observations imply that the methods used to construct the structural connectome, e.g. the choice of tractography method, parcellation resolution, and any subsequent thresholding, can affect all downstream results and limit the landscape of realizable staging patterns.

### 4.2 The graph Laplacian weight and the characteristics of growth versus diffusion

The graph Laplacian plays a distinctive role in reaction-diffusion models of prion-like propagation such as (2); it provides the mechanism for the transport of misfolded proteins over the brain’s structural network. As such, the choice of graph Laplacian weights has varied in the literature; particular weight selections are often motivated by a biological argument or an appeal to data fit. Overall, our results suggest that, while the choice of weights often has a minimal impact on the number of observable staging patterns of interest 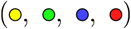, they did affect the prevalence of those patterns; in particular, the diffusive weights (DW) scheme tended to favor the expression of the progressive Braak 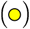 staging pattern over the options of Length-free weight (LW) and ballsitic weights (BW).

In the case of *τ*P seed staging, LW and BW showed significantly more sensitivity, in the observed staging, than the DW scheme as ln(*β*) drew nearer to zero. The staging sensitivity to the weighting scheme can be illustrated by computing the standard deviation of the staging time series. Plots of these standard deviations, for *τ*P seeding on the Scale-500 deterministic connectomes (see Figure 3), are shown in Figure 10. Regions, in Figure 3, where the staging is sensitive to small perturbations in *β* or *T* are precisely those where the standard deviation of the staging time series is low. This implies that the observed sensitivity is a *race condition* where the differences in threshold-attainment time are nearly identical.

**Figure 10:**
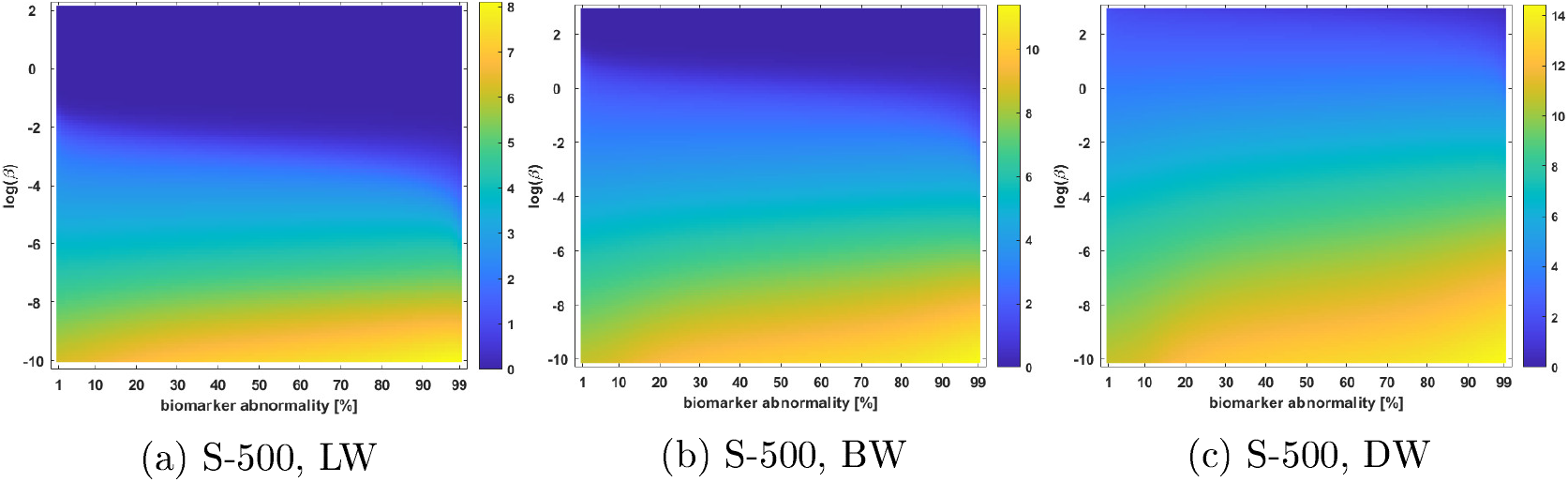
Standard deviation of the staging time series for *τ*P seeds on the high-resolution (S-500) deterministic connectome of Figure 3; all weighting schemes.

As length plays a more dominant role in the weighting scheme, the standard deviation of the staging time series increases, staging patterns are no longer sensitive to small perturbations and *τ*P seed staging becomes pronounced. However, it is not currently known if *τ*P seeds should follow a clear hierarchical progression though the appearance of *τ*P seeds have been shown to proceed NFT pathology [8]. In the case of NFT staging, the DW scheme generally promoted the progressive Braak pathway NFT staging pattern 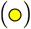 across all connectomes. This pattern, however, was not the SUVR-suggested 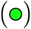 pattern according to our analysis. It is worth noting that the patient data necessary to achieve more statistically significant results, for latter stages, was not present (see Methods, Data preparation and Methods, Identifying additional computational staging patterns of interest). Moreover, as we mentioned previously, the II→III versus III→II paradigm, differentiating the progressive staging 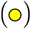 from the SUVR-suggested staging 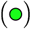, may be related to the prevalence of seeds 8 in the posterior parahippocampus at histopathological Braak classification stage II. Nevertheless, our results suggest that direct comparisons to flortaucipir data, at least for the ROI we considered, may be best conducted (see Figures f and f) with the BW scheme and on high resolution connectomes created with deterministic streamlining and with a choice of mathematical parameters given by −2 ≲ ln(*β*) ≲ 1.

Conversely, if a progressive Braak pattern 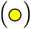 is desirable, this can most reliably be achieved, in general, on high resolution connectomes with the DW scheme, sufficiently dominant growth (i.e. *β* sufficiently small) and a typical biomarker threshold of at least 50%. As a point of practice, this should be checked with a braid surface analysis as we did observe that thresholding methods can perturb this trend on the probabilistic connectomes that we considered. For example, Figure shows that the density filter threshold favors the LW and BW schemes over the DW scheme for the progressive staging 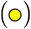; likewise, Figure shows that the high-salience scheme more readily expresses the progressive staging 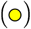 on lower-resolution connectomes.

### 4.3 Conclusion

Investigations of *τ*P staging have been carried out histopathologically [21, 22, 9] using SUVR [23, 10] and in fitting models of spreading to SUVR data [1, 2, 7]. As computational power is readily available, the prion-like hypothesis has enabled a new generation of mathematical models facilitating the latter types of studies. However, it is currently not known how various model selection choice may promote, or limit, the expression of particular staging patterns. We have found that every aspect of model selection can alter the overall progression of *τ*P staging. This includes parcellation resolution; tractography method; choice of thresholding method; choice of threshold; graph Laplacian weights; and the mathematical model parameters. We have shown that the the generalized staging problem couples the global topology of the structural connectome to the mathematical model governing the local dynamics of a biomarker evolving on that network and that comprehensive model selection cannot, in general, be ensured from a decoupled perspective. We have introduced braid diagrams and braid surfaces as a means to investigate the complex staging landscape, and changes thereto, for this coupled problem and to observe the implications of model selection choices.

The generalized staging problem in AD reveals the need for more experimental data and poses new theoretical questions. We propose that further study into the progression patterns of *τ*P seed staging, extending the results of [8], could promote a more reliable selection of parameter ranges for the balance of growth and diffusion (*β*) for *τ*P seeds and, due to the consistency we observed, for *τ*P NFT. Moreover, an increased number of SUVR data for patients in the latter Braak stages would help to belay uncertainty in determining the progression patterns suggested by data studies. Mathematically, the study of the coupling between the connectome topology, the weighted distances between ROIs, and the model parameters is a promising candidate for theoretical research.

## 5 Methods

### 5.1 Structural connectomes

Braid surfaces have been introduced to facilitate model selection, in the context of staging patterns developing on two sets of undirected, multi-resolution structural connectomes; both sets of connectomes were originally constructed from the data of participants in the HCP. All of the structural connectomes considered in this manuscript were constructed using the Lausanne multi-resolution atlas parcellation [27] with five levels of potential resolution; the coarsest scale (Scale-33), three intermediate scales (Scale-60, Scale-125, Scale-250), and a fine scale (Scale-500). The edges, at all scales, include information regarding the number of fibers (*n*_*ij*_) and fiber length (*ℓ*_*ij*_) associated to each edge (*e*_*ij*_) connecting region *i* to region *j*. The first set of connectomes (see Fig 11) were constructed using MRtrix [28, 29] and a deterministic streamlining with 20,000 streamlines and randomized seeding. These connectomes span five scales; the lowest and highest scales are shown in Figure 11. These connectomes are publicly available, as the dataset named ‘Full set, 426 brains, 20,000, streamlines’, from [24].

**Figure 11:**
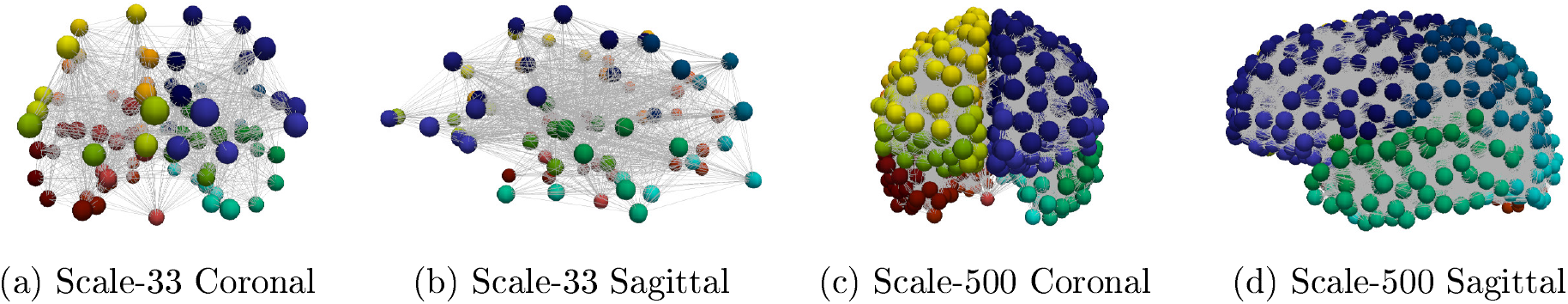
Lowest (left) and highest (right) connectome resolutions. Node colors signify the 83 disjoint anatomical parcellation regions.

The second set of connectomes were constructed for this study in order to assess degrees of certitude regarding observations for the deterministic connectomes mentioned above. The Connectome Mapping Toolkit [27] was used to parcellate a high resolution MNI reference template; the FSL [34, 35, 36] PROBTRACKX algorithm was employed, for the probablistic tractography, with 10,000 streamlines per voxel. The sparsity of the resulting connectivity matrices was low (approximately 7-12%); these matrices are commonly thresholded before use in computational models. To study the effect of thresholding, we considered five different thresholding techniques in our comparative analysis. The thresholding techniques used are summarized in Table 4 and a description of the method can be found in the corresponding citation. The naive thresholding method removes edge *e*_*ij*_ if the corresponding connectivity coefficient (*n*_*ij*_) is below a prescribed threshold value.

For comparison, we have constructed connectomes for the lowest (Scale-33, 50 patients) and highest scale (Scale-500, 25 patients) resolutions of the Lausanne multiresolution atlas. The braid surface source code, and the full set of thresholded connectomes, used in this study is available online [37].

### 5.2 A staging for *τ*P seeds on structural connectomes

We have discussed several hierarchical staging patterns for *τ*P progression in AD. As our mathematical model accounts for both *τ*P seeds and *τ*P NFT, we selected a six-stage model that has been related to both quantities [8]. These six regions, along the Braak pathway, are enumerated in Table 5.

**Table 5:**
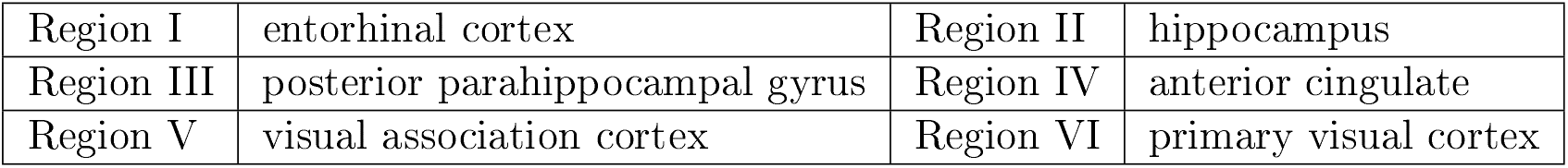
Anatomical staging regions for *τ*P seeds as reported in [8].

The regions in Table 5 can be mapped to anatomical ROI of the Lausanne atlas used in the construction of the structural connectomes for this study. We therefore adapt the stages of Table to the Lausanne atlas by considering the staging process of Table 2. The primary difference is that stages V and VI, of Table 5, are combined into a single terminal stage and approximated by the presence of deposition in the cuneus, pericalcarine cortex, lateral occipital cortex, and lingual gyrus.

### 5.3 Staging sequences for *τ*P pathology on computational connectomes

In order to assess the implications of various model parameters on observed computational staging patterns, thus facilitating model selection, a choice of preferable staging patterns must be identified. We select five collections of nodes, Ω_*i*_ for *i* = 1, 2, … , 5, such that Ω_*k*_ is the set of nodes of the computational connectomes whose anatomical ROI labels are given by the *k*^th^ stage of Table 2; thus Ω_1_ contains all nodes labeled as belonging to the left and right entorhinal cortices, etc. The staging regions, Ω_*i*_, for the coarsest (top row) and finest (bottom row) connectome resolutions are shown in Figure 12

**Figure 12:**
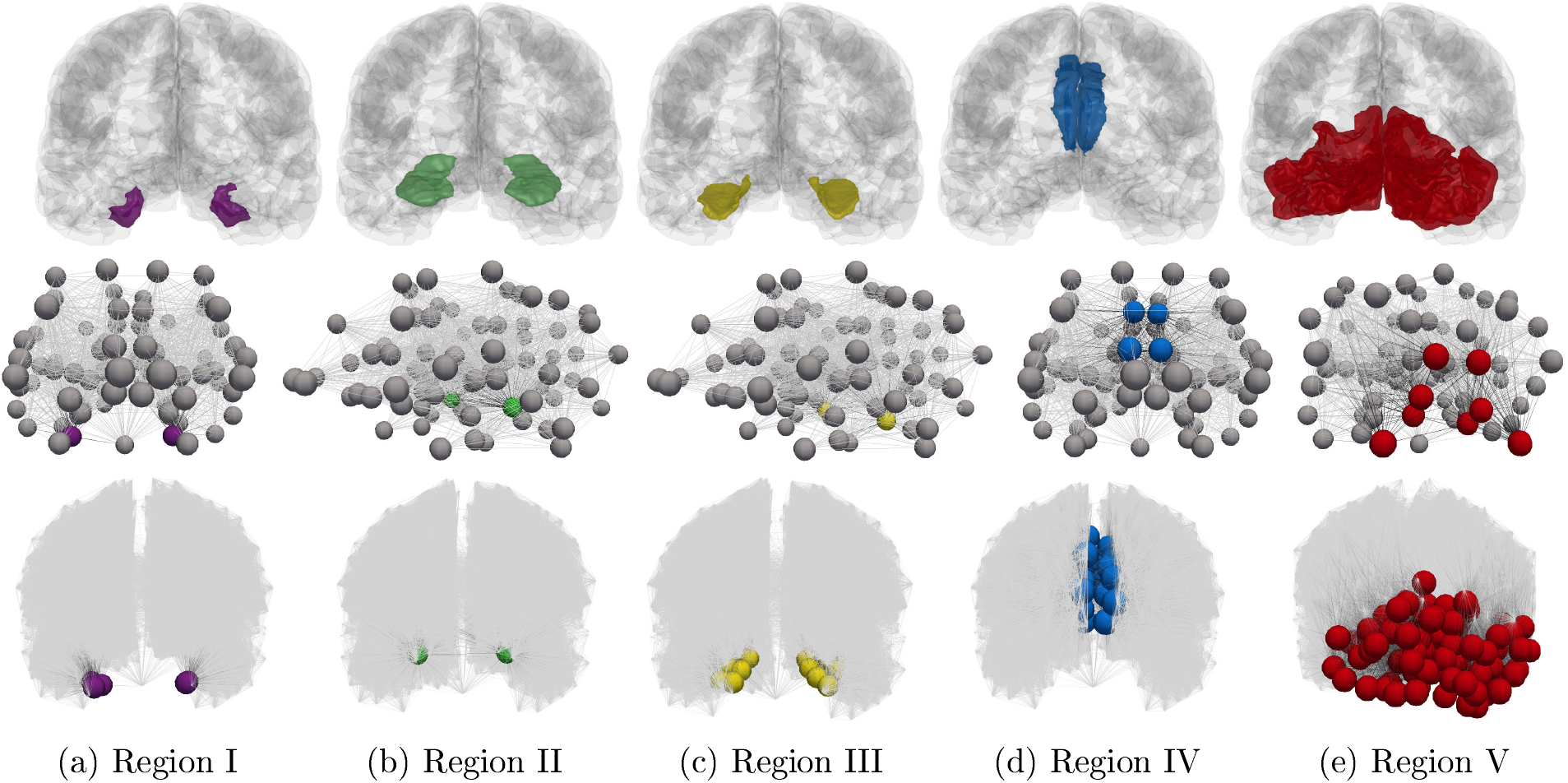
Connectome Braak staging regions, Ω_*i*_. Anatomical regions in a glass brain (top). Coarsest connectome resolution (Scale-33 parcellation, middle) and finest resolution (Scale-500 parcellation, bottom, additional nodes suppressed) connectomes.

The work of [8] assessed *τ*P seed staging along the Braak tau pathway and found that tau seeding precedes the presence of NFT pathology. A clear candidate for both *τ*P seed and NFT staging is therefore the progressive regional staging I → II → III → IV → V. To assess other potential staging sequences we used flortaucipir (AV-1451) data from the Alzheimer’s disease neuroimaging initiative (ADNI); the data was fully preprocessed at the Helen Wills Neuroscience Institute at the University of California, Berkeley and the compiled results are freely available through the ADNI website. The preprocessed dataset contains 1,184 records; each record corresponds to a subject structural MRI scan and tau PET scan and the preprocessing steps are explained in the companion document available through the ADNI website.

#### 5.3.1 Data preparation

The Berkeley preprocessing pipeline is as follows: each patient T1 MRI was segmented using FreeSufer 7.1.1; each subject’s normalized-intensity flortaucipir scan was co-registered to their bias-corrected T1 image; partial volume effects were corrected for using the geometric transfer matrix approach [38, 39]; and mean flortaucipir uptake is reported, in the dataset, for each region of the FreeSurfer segmentation. We further normalized each subject’s regional SUVR score using the inferior cerebellar gray matter as the reference region, as suggested in the dataset documentation.

We proceeded to create two datasets using this preprocessed dataset. For the first set, we computed the volume-weighted SUVR average, for each patient record: over the whole brain to compute a global score (GS); and for each region of Table 2. Thus, each ADNI patient visit record was assigned a global volume-averaged SUVR score along with five volume-averaged regional scores corresponding to the regions of Table 2. We refer to this dataset as the ‘base dataset’.

We then created a second dataset we refer to as the ‘partitioned dataset’ by following the methodology of [23]. An overview of the creation process follows First, the GS was used as a decision variable, in a conditional inference tree (CIT) algorithm. Second, the CIT algorithm was used to compare the GS to the BraakV regional volume-weighted SUVR; producing a GS threshold. This GS threshold was used to separate patient records into two groups who differed significantly based on their BraakV score. The BraakV patient records were then moved, out of the full dataset, into a new data file. This process was then repeated; once again applying the CIT algorithm and using the GS as a decision variable, but now comparing to the BraakIV regional volume-weight SUVR score. Thus creating a BraakIV group of patient records, and so on. This process resulted in five separate files, i.e. the partitioned dataset, corresponding to a patient record classification according to the regions of Table 2. The thresholds produced by the CIT algorithm can be seen in Table 6. Our application of CITs follows the identical approach of previous authors [23], in a similar SUVR study, to categorize each patient record into the discrete groups corresponding to a pre-defined set of regional stages.

**Table 6:** Conditional inference-based partitioning of ADNI flortaucipir data into the stages of Table 2

| Stage group classification | Volume-averaged global SUVR (GS) | Subject records in group |
| --- | --- | --- |
| I | $1.266 \leq \text{GS}$ | 36 |
| II | $1.266 < \text{GS} \leq 1.392$ | 248 |
| III | $1.392 < \text{GS} \leq 1.52$ | 443 |
| IV | $1.52 < \text{GS} \leq 2.365$ | 416 |
| V | $\text{GS} > 2.365$ | 41 |

#### 5.3.2 Identifying additional computational staging patterns of interest

Our next goal was to see if ADNI flortaucipir data may suggest alternatives to the progressive I → II → III → IV → V (computational) Braak-like sequence; in particular for NFT due to the binding of flortaucipir to paired helical filaments. Our objective here is not to rigorously study hierarchical SUVR behavior as in [10]. Rather, our goal is to conduct a simple inquiry as to whether the ADNI data may suggest alternative staging patterns of potential interest for the computational staging problem and model selection; especially as it pertains to observed NFT staging.

Our first investigation used the partitioned dataset. First, we checked that the CIT algorithm separated the data in such a way that all of regional mean SUVR values, across all possible group-region pairings, were significantly (*p* < 0.05) different, using a Welch’s t-test; this was the case, as expected by an application of CIT-based separation. Next, we computed the Pearson correlation between each stage group’s primary stage and all other stages within that group; the results are reported in Table 7.

**Table 7:**
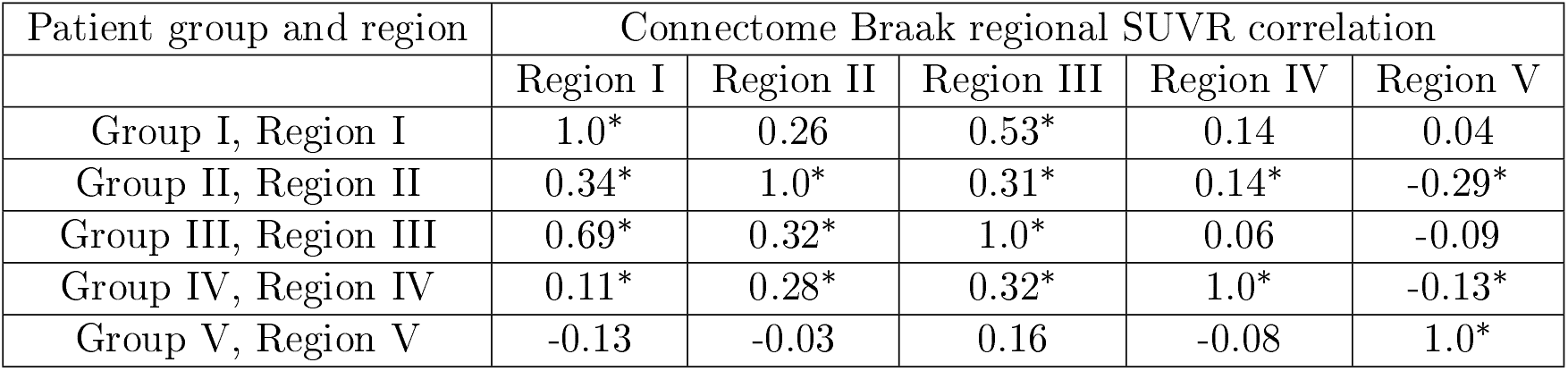
Pearson correlation between a stage group’s primary region and all other regions within the group. A ∗ denotes a statistically significant correlation (*p* < 0.05) between regions.

Assuming that overt NFT pathology originates in region I, Table 7 suggests that staging sequences with I → III may also be of interest for NFT. Assuming that III proceeds I, the next suggested progression would be III → II. This observation suggests that sequences starting with the prefix I → III → II could be of interest alongside the more progressive stage prefix I → II → III. Finally, likely to to the sparsity of patient records assigned to Group V, none of the correlations of Group V were significant.

Our next investigation uses the base dataset. The primary motivation was to ensure that patient gender and age were not confounding our partitioning based strategy. Starting with the base dataset, we cross-referenced ADNI patient data and augmented the base set with the age, at the time of scan, in addition to the patient’s gender. We then used regression models covarying for age and gender. We compared each regional volume-averaged SUVR score to that of the other regions; the results are shown in Table 8. Evidence for influence was evaluated using standardized factor scores when the influence was significant. Region III was most affected by Region I (*R*^2^ = 0.775, *B* = 0.613, *p* < 0.001); further supporting interest in the I → III staging prefix. Region II was also most influenced by Region I (*R*^2^ = 0.486, *B* = 0.486, *p* < 0.001) but at a lower value of both *B* and *R*^2^. Removing Region I from the model decreased the variational fidelity (*R*^2^) of the Region III model by 24.8% and the Region II model fidelity by 13.6%; in the latter case, the Region II score was most affected by the Region III score (*R*^2^ = 0.42, *B* = 0.5). Supposing that pathology begins in the entorhinal cortex (Region I), these observations suggest, again, that a I → III → II prefix is also of interest for evaluating computational staging.

**Table 8:** Regression models, covarying for age and gender, of regional influence

| Regression model | Influential model region(s) | Model $R^2$ | $B$ | $p$-value |
| --- | --- | --- | --- | --- |
| Region II | Region I | 0.486 | 0.49 | $p < 0.001$ |
| | Region III | 0.42 | 0.5 | $p < 0.001$ |
| Region III | Region I | 0.78 | 0.61 | $p < 0.001$ |
| | Region IV | 0.58 | 0.39 | $p < 0.001$ |
| | Region II | | 0.36 | $p < 0.001$ |
| Region IV | Region III | 0.43 | 0.75 | $p < 0.001$ |
| Region V | Region III | 0.22 | 0.38 | $p < 0.001$ |

Next, we take a second look at the staging suffix options IV → V versus V → IV. To do so, we regressed the Region IV and Region V scores in terms of the first three regional scores, once more covarying for age and gender. The Region IV model identified Region III as the most influential (*R*^2^ = 0.43, *B* = 0.75, *p* < 0.001). The Region V model was only marginally explained by the data (*R*^2^ = 0.22), but the most influential factor was, again, the Region III score (*B* = 0.38, *p* < 0.001) but with a substantially lower influence than in the Region IV model. These latter observations suggest that a IV → V may be more desirable than a V → IV suffix. We have therefore corroborated, and extended, the conclusions suggested by the partitioning strategy (Table 7).

Next, we corroborate our previous SUVR findings by following a similar approach to [10] and considering Z-scores, computed from the base dataset, for the regions defined in Table 2. First, all of the regional mean SUVR scores were individually standardized. Similar to [10], we considered a particular region to be ‘involved’ in a fixed patient record if that region’s Z score satisfied *Z* ≥ 2. We then selected all patient records with an involved Region I (entorhinal cortex). Continuing within this subset of data, we counted the frequency of all other involved regions and computed their mean Z-score. The results are reported in Table 9. Once more, we see that progression sequences starting with I → III → II may also be of computational interest and, further, that the staging suffix IV → V appears more preferable, for SUVR, than the V → IV alternative.

**Table 9:** Spreading order of *τ*P (SUVR) for Region-I involved patient records

| Region | % Involved | Z |
| --- | --- | --- |
| Region I | 100.0 | 3.24 |
| Region III | 55.1 | 2.55 |
| Region II | 46.94 | 1.81 |
| Region IV | 22.45 | 1.43 |
| Region V | 22.45 | 1.17 |

Finally, we compared the ADNI data findings, above, to the results for *τ*P seeds reported in [8] where Braak II brains (classified post-mortem) also showed a slight average increase of *τ*P seeds in physiological region III (i.e. region III, Table 5) over that of physiological region II; further suggesting that a computational I → III → II pattern may be of interest for *τ*P seeds. If one considers post-mortem Braak classification as a pseudo-time then, from [8, Fig. 3], it is tempting to suggest a computational I → III → II → IV → V pattern of interest for *τ*P seeds. However, the differential between seeding levels in the latter stages is slight. In summary ADNI SUVR data, and results from [8], suggest a broader set, beyond the choice of a strictly ascending progression, of potentially interesting computational staging patterns for the choice of regions in Table 2. These staging patterns are reported in Table 10. We note that either more late stage data (i.e. Group V, Table 6), a finer SUVR staging sequence, such as that of [10], or both is needed to ascertain a higher degree of certainty in the latter *τ*P computational staging patterns for AD.

**Table 10:** Computational staging patterns of potential interest for *τ*P seeds and NFT. * Progressive Braak staging, ^†^ Suggested by SUVR data, ^‡^ Additional connectome computational stagings of potential interest pertaining to uncertain *τ*P seed staging prefixes.

| Computational staging patterns (see Table 2) |  |  |
| --- | --- | --- |
| Canonical late staging | $I \rightarrow II \rightarrow III \rightarrow IV \rightarrow V^*$ | $I \rightarrow III \rightarrow II \rightarrow IV \rightarrow V^\dagger$ |
| Uncertain late staging | $I \rightarrow II \rightarrow III \rightarrow V \rightarrow IV^\ddagger$ | $I \rightarrow III \rightarrow II \rightarrow V \rightarrow IV^\ddagger$ |

## Acknowledgement

The work of A. Goriely was supported by the Engineering and Physical Sciences Research Council grant EP/R020205/1. The work of P. Putra was partially supported by the Engineering and Physical Sciences Research Council grant EP/R020205/1 to AG. The work of T. Thompson was supported partially the John Fell Oxford University Press Research Fund grant 000872 (project code BKD00160) to TT, and partially by the Engineering and Physical Sciences Research Council grant EP/R020205/1 to AG. The work of P. Chaggar was supported by funding from the Engineering and Physical Sciences Research Council grant EP/L016044/1 and Roche.

Data used in the preparation of this article were obtained from the Alzheimer’s Disease Neuroimaging Initiative (ADNI) database (adni.loni.use.edu). The ADNI was launched in 2003 as a public-private partnership, led by Principal Investigator Michael W. Weiner,MD. The primary goal of ADNI has been to test whether serial magnetic resonance imaging (MRI), positron emission tomography (PET), other biological markers, and clinical and neuropsychological assessment can be combined to measure the progression of mild cognitive impairment (MCI) and early Alzheimer’s disease (AD). For up-to-date information, see www.adni-info.org. A complete listing of ADNI investigators can be found at http://adni.loni.use.edu/wp-eontent/uploads/how_to_apply/ADNI_Aeknowledgement_List.pdf

## Technical Terms

### Diffusion-reaction

A diffusion-reaction model refers to a system of semi-linear parabolic partial differential equations describing the evolution of quantities **p** (typically chemical concentrations) throughout a medium. A typical form for these equations is *∂*_*t*_**p**(**x**, *t*) = ∇·(**K**∇**p**(**x**, *t*))+**R**(**p**(**x**, *t*)) where **x** ∈ ℝ^3^ represents space, *t* is time, **K** is a diffusion tensor and **R**(**p**(**x**, *t*)) is a nonlinear function of its argument. This type of system forms the basis of discussion in the section ‘A connectome diffusion-reaction model of *τ*P proteopathy’ (Theory and model).

### Grap Laplacian

The graph Laplacian, otherwise known as the Laplacian matrix, is a matrix representation of a weighted or unweighted network; it can be used to study many properties of the network. In the case of an undirected network, the graph Laplacian is often interpreted as the discrete analogue of the diffusion operator.

### Model selection

The term model selection refers to the process of arriving at the set of features that determine the model used to study a process of interest.

### Prion-like

Prions are infectious proteins that can exhibit aberrant conformational properties. Normal forms of prion proteins can themselves become misfolded in the presence of another already-misfolded prion variant; leading to so-called prion diseases. The term ‘prion-like’ refers to proteins, other than prion proteins, that replicate via self-propagating conformational abnormalities; assembling into filaments that are infectious through their ability to grow by recruiting the soluble form of the protein [40].

### Simplicial mesh

A mesh is typically used to refer to a discretization of an, often complex, geometric domain of interest. A simplicial mesh is a mesh whose elements are triangles or tetrahedrons and is a widely used option in finite element, or finite volume, discretizations of partial differential equations.

### Transversely anisotropic

A transversely anisotropic material is one that exhibits symmetry, with respect to some material property, about an axis that is orthogonal to a plane in which that property is isotropic (i.e. the same in all directions). In our case, the material property of interest is (water) diffusion in the brain.

## Supplementary Information Sl: Diffusion reaction systems and the conservative graph Laplacian

### A continuous diffusion-reaction model

Our present focus is on non-linear diffusion-reaction models. The typical scalar is form for such models is

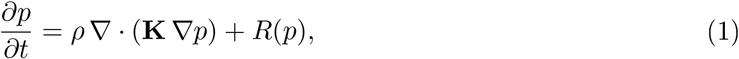

where *p* is the concentration of misfolded *τ*P, *R*(*p*) represents the non-linear reaction term for the model, **K** is a (symmetric) matrix diffusion tensor and *ρ* is an effective diffusion constant; when the diffusion is isotropic, **K** is a multiple of the identity matrix and ∇ · (**K** ∇*p*) = *ρ*Δ*p* is the usual continuous Laplacian operator. In the brain, axonal white-matter bundles heavily bias the diffusion of extracellular fluid; leading to a transversely anisotropic diffusion tensor [1] with general form

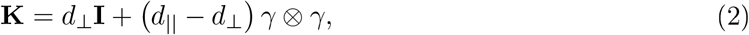

where **I** is the identity matrix, *d*_⊥_ ≪ *d*_∥_ and *γ* is a unit vector oriented along the fiber bundle. We now note two properties of (1). First, the form of the diffusive term, ∇ · (**K**∇*p*), in (1) is a direct result of Fick’s law of diffusion. Second, in the absence of any explicit mass exchange terms, such as production or clearance, or specific reaction terms, the model (1) conserves mass; a feature that should be consistent in a discretized version of the problem.

In the main manuscript, we study the generalized staging problem, on networks, of the simplest, single-species model for the prion-like propagation of *τ*P proteopathy which promotes both prionlike propagation and growth; this Fisher-Kolmogorov-Petrovsky-Piskunov (Fisher-KPP) model has been used in several previous investigations [1, 2, 3, 4, 5]. This model allows for a direct study of staging in the presence of prion-like growth (due to the reaction terms) and diffusive transport. The Fisher-KPP model is defined by setting *R*(*p*) = *αp*(1 − *p*) to obtain

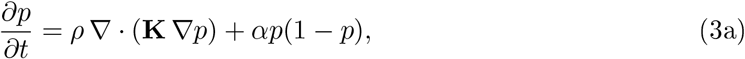

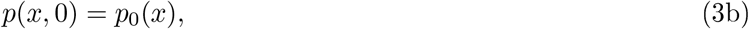

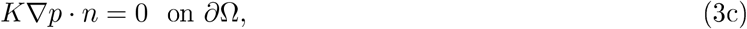

where 0 ≤ *α* determines the local growth rate, potentially varying smoothly in space, and *p*_0_(*x*) is the initial seeding of *τ*P. We will be particularly interested in the observed staging behavior, along the Braak pathway, of network discretizations of (3) with respect to the relative size of the parameters *ρ* and *α*; the Fisher-KPP model has two asymptotic regimes:

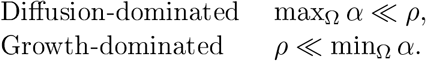

### Connectome discretizations of diffusion-reaction models

The problem now is to find a suitable discretizations of the spreading process by using the network structure of the brain. While some mouse models are sufficiently precise to establish directionality and hence build a directed connectome [6, 7], parallel evidence for directed Ifow within the human brain is lacking. Moreover, there is no available directional human structural connectome and it has been shown, theoretically, that directionality has very little appreciable impact on the overall spreading dynamics [5]. We therefore consider an undirected connected network *G* = (*V, E*) with |*V*| = *N* ≥ 3 nodes and |*E*| = *M* ≥ 2 edges. We further assume that the network is weighted with positive weights **W**_*ij*_ = **W**_*ji*_ at an edge joining node *i* to node *j*. Weights are assumed to be zero if two nodes are not connected, strictly positive if two nodes are connected and the graph is assumed to be free of self-loops (i.e. **W**_*ii*_ = 0) at each vertex. The weighting matrix **W** therefore defines an *N* × *N* symmetric *weighted adjacency matrix* on the undirected graph *G*. For the remainder of the discussion, we presume that the graph *G* represents a structural connectome of the human brain. That is, the vertices of *G* represent regions of interest (ROI), determined by a choice of parcellation, and the edges of *G* represent the white matter fiber bundle connectivity between the ROIs.

Within the brain, the prion-like propagation of misfolded proteins is dominated by its transport along the axonal fiber bundles. It is therefore natural to consider propagation along edges of the structural connectome graph, *G*, discussed above. To do so, we replace scalar fields, such as *p*, *∂*_*tp*_, *α*(*x*) etc, with regional (ROI) averages, denoted by *p*_*i*_, 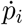, or *α*_*i*_ etc, and replace the continuous diffusion operator ∇ · (**K**∇*p*) with a graph Laplacian. This type of discretization strategy can be considered as motivated by, for instance, finite volume methods [8, 9] applied, in space, to (1) with a simple quadrature, such as the midpoint rule, for estimating the (spatial) integral of the non-linear term, *R*(*p*), on a mesh whose cell faces are constructed orthogonal to the fiber directions. A natural discretization of (1) is therefore

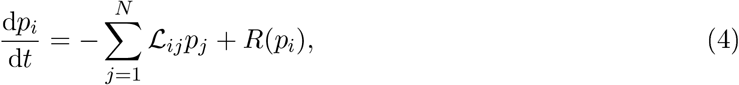

where *p*_*i*_ is the concentration at each ROI *i* and 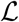 is the *N* × *N* graph Laplacian. The standard graph Laplacian has the form

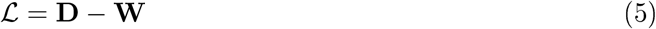

where **W** is a weighted adjacency matrix that codifies the connectivity between vertices of the graph *G* and **D** is the diagonal degree matrix defined by

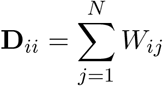

The graph Laplacian matrix can also be normalized, which we will discuss in the following section. Using the graph Laplacian, the full connectome graph discretization of (3) is

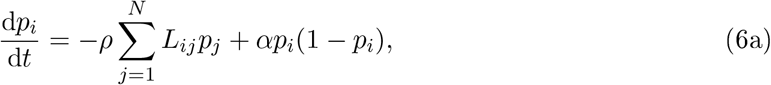

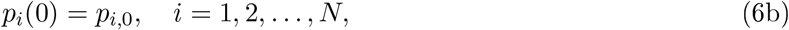

The continuous Neumann boundary condition (3c) is enforced, at the network level, by ensuring that the graph Laplacian robustly conserves mass; mass conserving forms of the graph Laplacians are the topic of the next section. We close by mentioning that (6) can be rescaled in time to produce the model considered in the primary manuscript; namely,

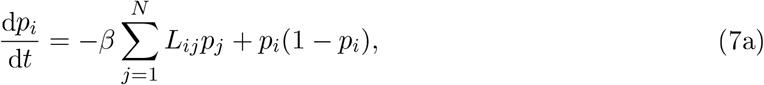

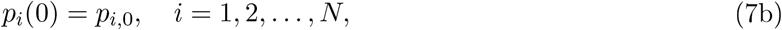

where *β* = *ρ/α*.

### The many graph Laplacians

In this section we consider the properties of mass conservation and Fick’s law with respect to the form of the graph Laplacian matrix 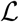. To motivate this investigation, we consider (7a) without the additional reaction term; that is,

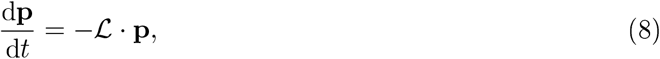

where **p** = (*p*_1_, … , *p*_*n*_) is the (column) vector of concentrations. First, assume that each ROI has the same volume; we will consider the case of varying volumes momentarily. Mass conservation, which we state as condition (C1) below, requires that

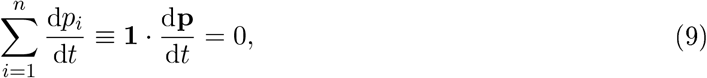

where **1** = (1, … , 1) is the one vector. Using (8), this condition implies

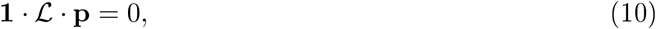

which must be true for all **p**. Hence we have the condition

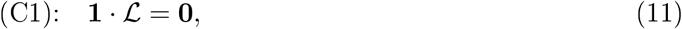

where **0** = (0, … , 0) is the null vector. Fick’s condition, which we state as (C2) below, states that in the absence of a concentration gradient, there is no transport; this statement is e uivalent to the condition that

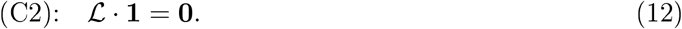

When the graph Laplacian is symmetric, conditions (C1) and (C2) are mathematically equivalent. In addition to (C1) and (C2) we also consider a robustness condition, stated as

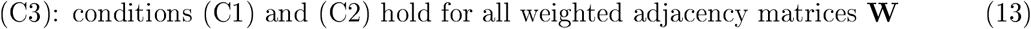

Condition (C3) states, in particular, that variations in the weighted adjacency matrix should not incur a loss of the fundamental physical principles reIfected by conditions (C1) and (C2). This is an important practical re uirement, as graph Laplacian weightings for structural connectomes are often derived from tractography algorithms, determining the number of fibers and their lengths, which may report different values when different software packages are used.

#### A family of Laplacians

The standard graph Laplacian (5) has been used in several modeling studies [3, 10, 11] of network neurodegeneration. Normalized forms of the graph Laplacian have also been used to study network spreading of proteopathy [12], the relationship between structural and functional connectivity [13], atrophy in alzheimer’s disease [14], and pathology in both supranuclear palsy [15], and Parkinson’s disease [16]. The *normalized graph Laplacian* used in these works is

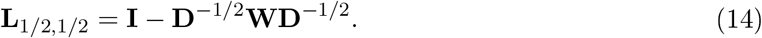

Other possible Laplacian matrices can be defined and have been used in various studies such as spectral clustering, including

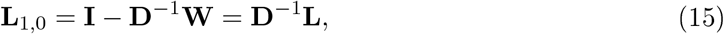

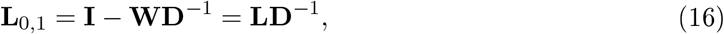

which motivates the introduction of the family

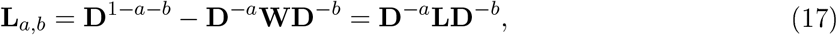

with *a, b* ∈ [0, 1] and *a* + *b* ≤ 1. The standard graph Laplacian corresponds to the choice *a* = *b* = 0. That is, **L** = **L**_0,0_ and we note that since the network is connected, the degree of each ROI (node) is strictly positive and the inverse of **D** is well defined. We can now state the main result of this section

##### 1. Proposition

*Suppose G* = (*V, E*) *is a connected undirected network with weighted adjacency matrix* **W**. *Then, the standard graph Laplacian* **L**_0,0_ = **L** *is the only member of the family* (17) *that simultaneously satisfies conditions (C1), (C2) and (C3)*.

*Proof*. From the identities

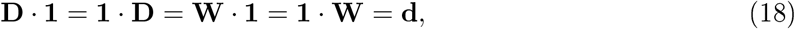

It follows that

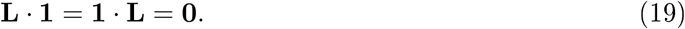

Hence the standard Laplacian **L** satisfies conditions (C1) and (C2). Condition (C3) is satisfied by the fact that these relationships do not depend on the form of **W**. Hence, changes in the weight will not affect either condition.

Next, we show that **L**_0,0_ is the only member of the family (17) satisfying all of (C1), (C2) and (C3). Let *a, b* ∈ [0, 1], with *a* + *b* ≤ 1, be arbitrary but fixed with at least one of *a* > 0 or *b* > 0.

First, consider conditions (C2) and (C3). From (17), and (C2) we have

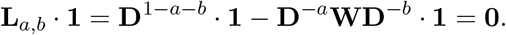

The *i*^th^ component of the above equation is

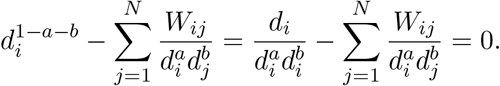

Multiplying through by 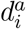 and using the definition of **D** in terms of **W** yield

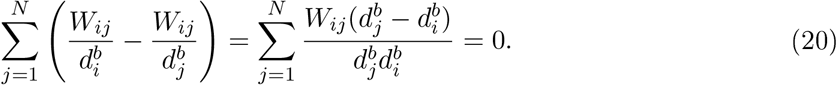

Since this identity must be respected for all undirected connected graphs (condition (C3)), it must be independent of **W**, hence we must have

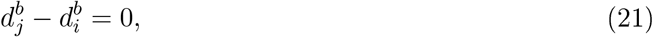

for all pairs (*i, j*). This condition is satisfied by either *b* = 0 or *d*_*i*_ = *d*_*j*_. If *d*_*i*_ = *d*_*j*_ for all ROIs (nodes) and since there are at least 3 ROIs (nodes) in the network, one can change the weight of a single edge connected to one of the ROI, say *i*, but not the other by adding an arbitrarily small amount 0 < *ϵ* ≪ 1 to that weight, with the effect of changing *d*_*i*_ but not *d*_*j*_. Hence, the e uality *d*_*i*_ = *d*_*j*_ cannot hold under the robustness assumption and we conclude that *b* = 0 is the only condition for which (C2) and (C3) hold simultaneously.

Next we consider the conditions (C1) and (C3). From (17), and (C1) we have

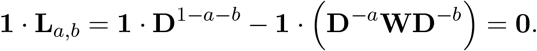

The *j*^th^ component of the column vector corresponding to the above equation states that

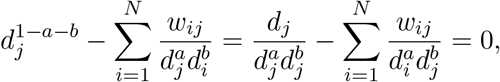

must hold identically. Multiplying through by the common term 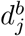 and using the symmetry of the adjacency matrix yield

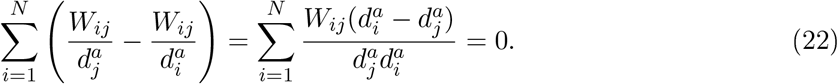

As in the previous case, the robustness condition (C3) implies that

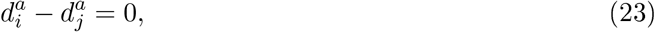

for all pairs (*i, j*) and we conclude that mass conservation and robustness imply that *a* = 0. Taken together conditions (C1), (C2) and (C3) imply that *a* = *b* = 0 must follow.

### Graph Laplacian correction for varying volumes

If the ROIs (nodes) have different volumes ***ν*** = (*ν*_1_, … , *ν*_*n*_), then the condition for the conservation of mass (C1) has to be modified. Indeed the total mass is now *P* = ***ν*** · **p**. nforcing 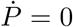 in (8) implies

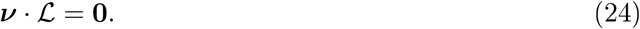

This condition can easily be achieved by choosing

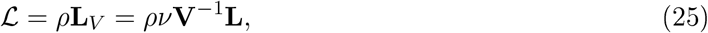

where **V** = diag(***ν***) and *ν* is a characteristic volume (average volume for instance). We note that this modified graph Laplacian is not symmetric, but that Fick’s condition is still satisfied (**L**_*V*_ · **1** = **0**) since diffusion takes place when a concentration gradient is established, independently of the ROIs volume. We also note that the multiplication of the standard Laplacian on the left by a diagonal matrix has been used to define regions of vulnerability [6, 7], hence assuming that diffusion takes place differently in different ROIs. Mathematically, it is the same operation but its interpretation in term of volumes or vulnerability is different and corresponds to different modeling choices (i.e. if we insist on mass conservation then mass should be conserved and interpreting **V**^−1^ as vulnerability precludes mass conservation).

## Supplementary Information S2: Additional staging results

### Additional results, deterministic streamlined connectome staging

**Figure 1:**
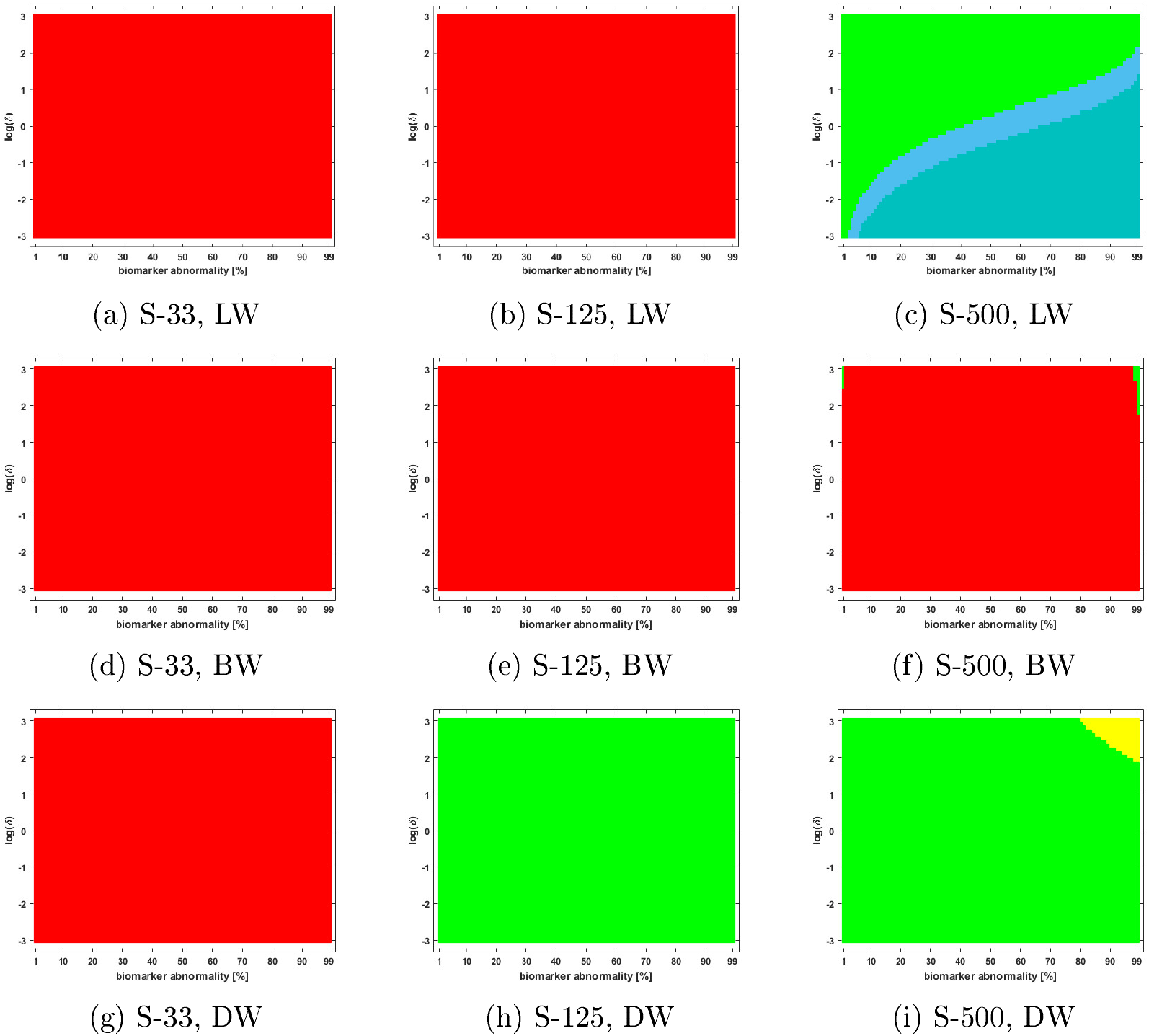
Braid surfaces, observed computational *τ*P NFT staging with deterministic streamlined connectomes; diffusion dominated regime (ln(*β*) = 2). Length-free (top), ballistic (middle) and diffusive (bottom) weighting schemes. The x-axis determines the biomarker abnormality threshold 1% < *T* ≤ 100% and the y-axis corresponds to NFT aggregation rate (*δ*) with −3 ≤ ln(*δ*) ≤ 3.

**Figure 2:**
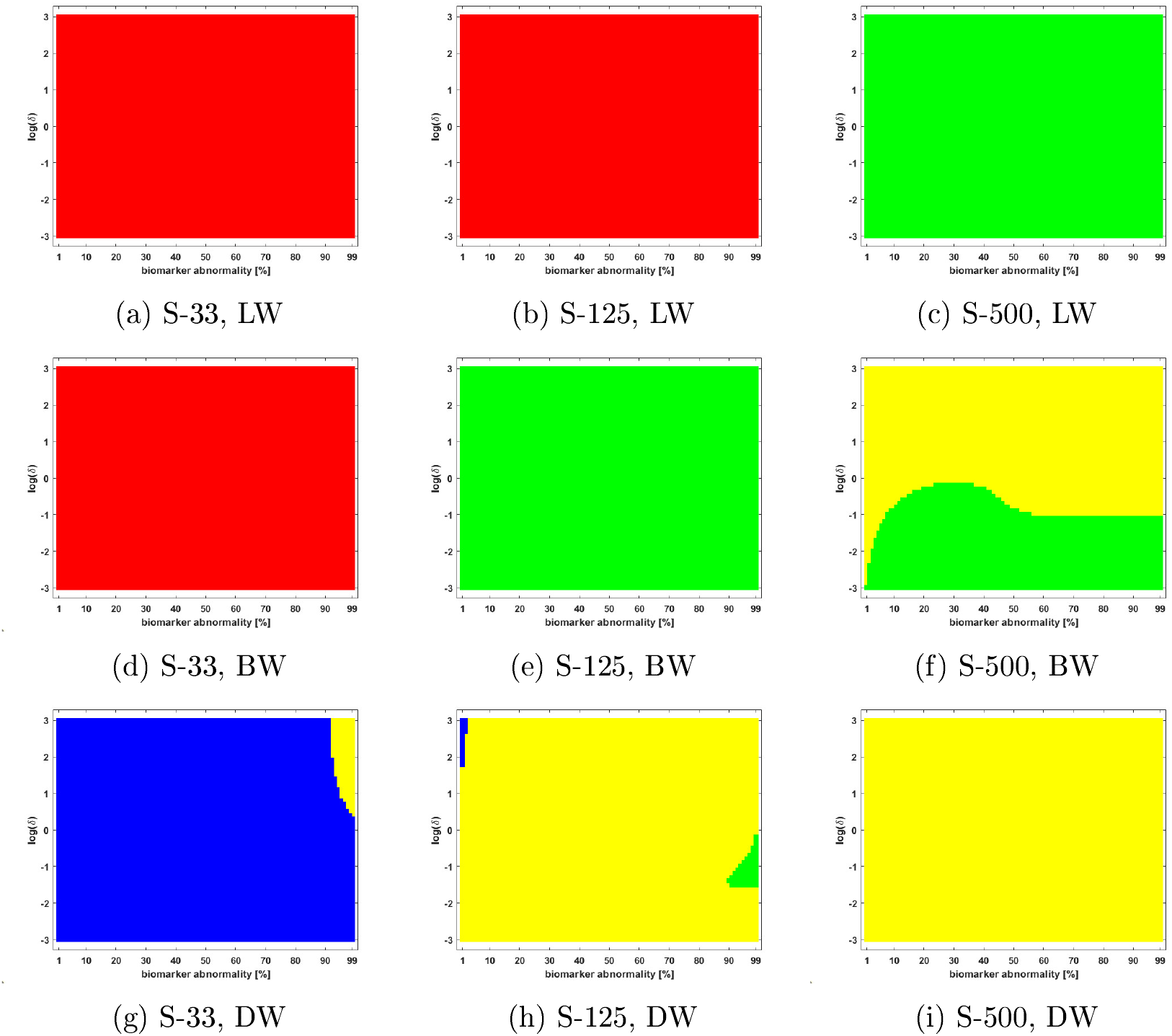
Braid surfaces, observed computational *τ*P NFT staging with deterministic streamlined connectomes; growth dominated regime (ln(*β*) = −3). Length-free (top), ballistic (middle) and diffusive (bottom) weighting schemes. The x-axis determines the biomarker abnormality threshold 1% < *T* ≤ 100% and the y-axis corresponds to NFT aggregation rate (*δ*) with −3 ≤ ln(*δ*) ≤ 3.

### Additional results, probabilistic streamlined connectome staging

**Figure 3:**
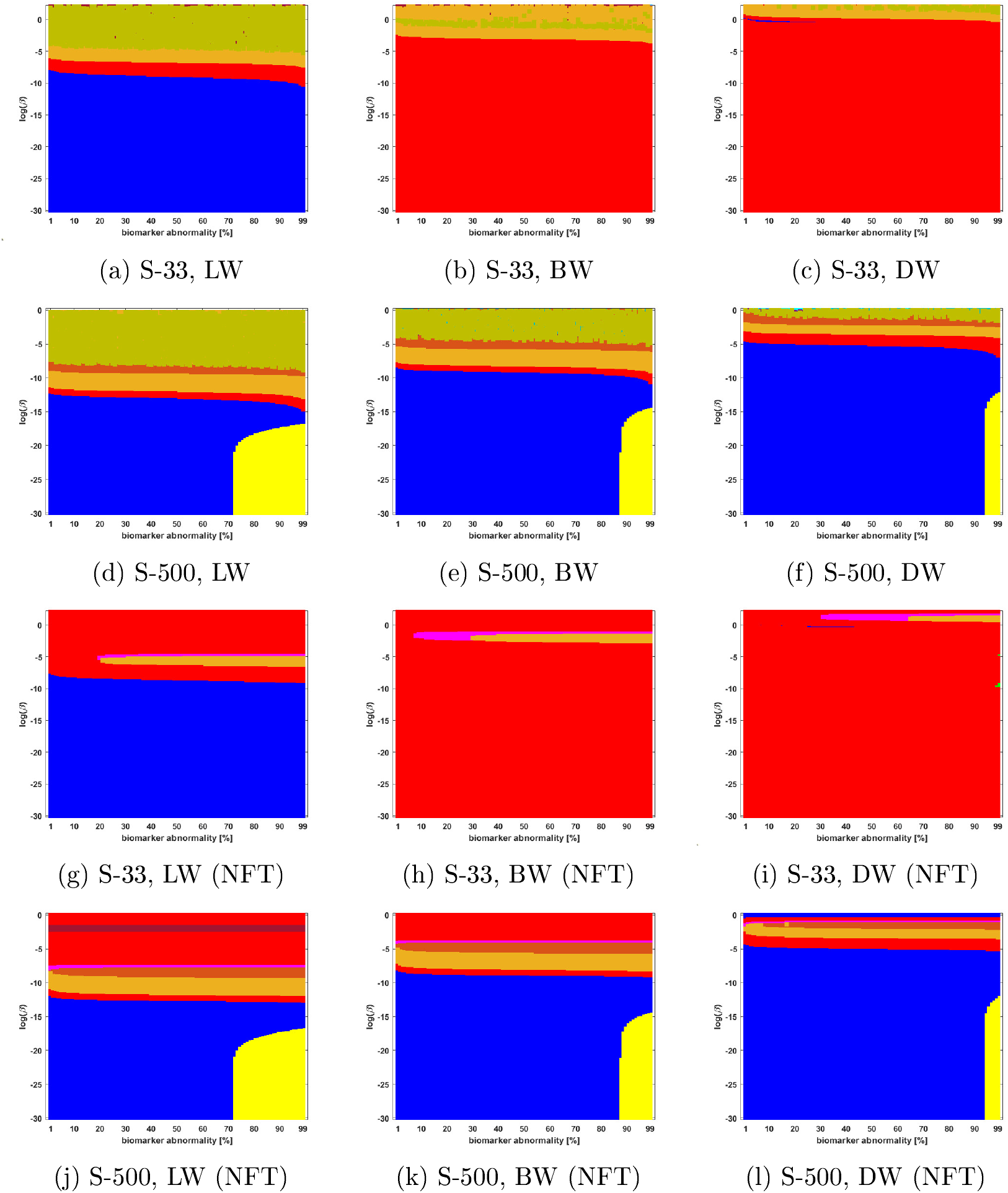
Observed computational (probabilistic) connectome *τ*P seed staging (top two rows) and *τ*P NFT staging (bottom two rows). Density filter thresholding at a threshold of 8 × 10^−1^ with biomarker abnormality 1% ≤ *T* ≤ 100% (x-axis) and −30 ≤ ln(*β*) ≤ 0 (y-axis)

**Figure 4:**
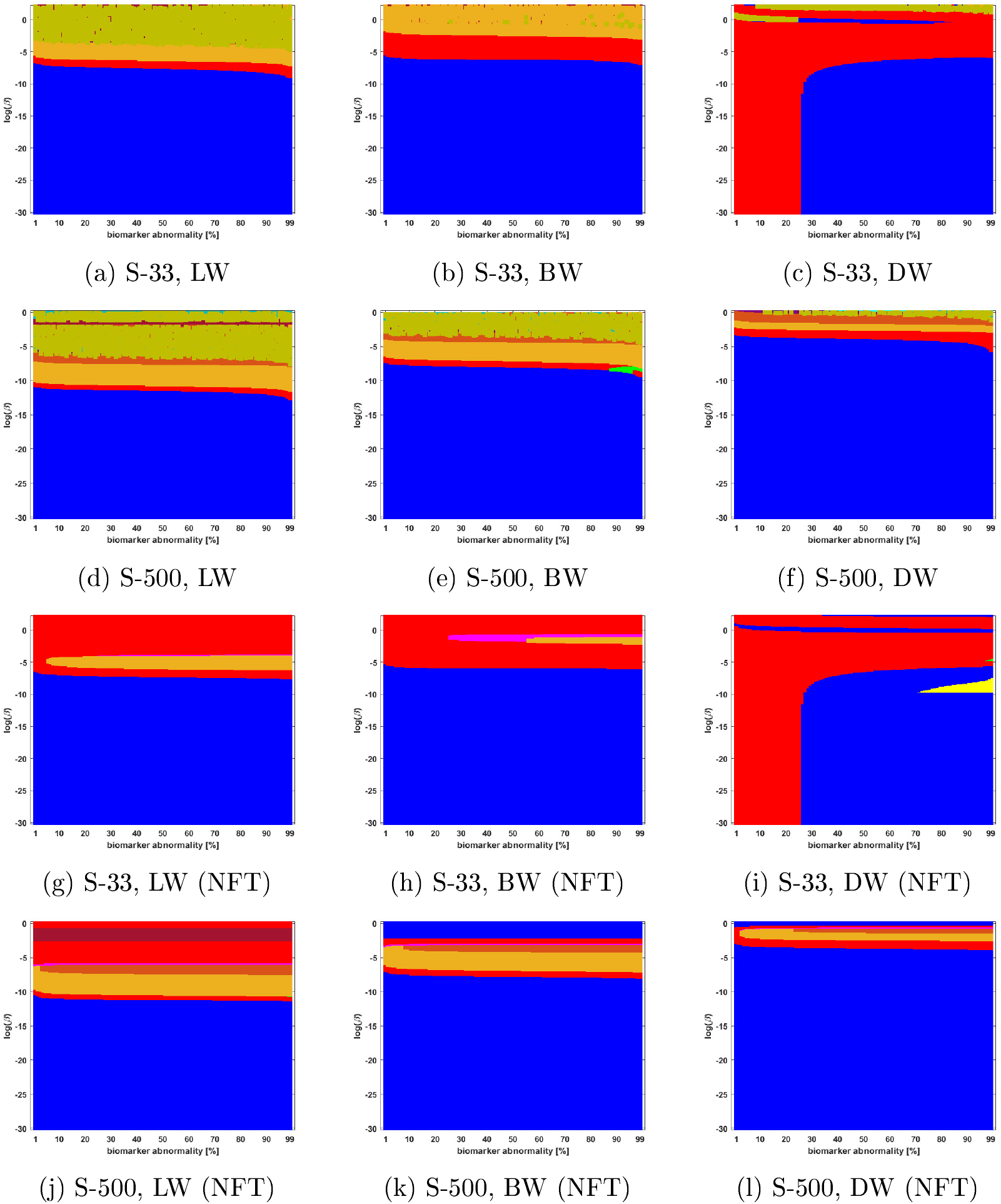
Observed computational (probabilistic) connectome *τ*P seed staging (top two rows) and *τ*P NFT staging (bottom two rows). High salience skeleton at a threshold of 5 × 10^−4^ with biomarker abnormality 1% ≤ *T* ≤ 100% (x-axis) and −30 ≤ ln(*β*) ≤ 0 (y-axis)

**Figure 5:**
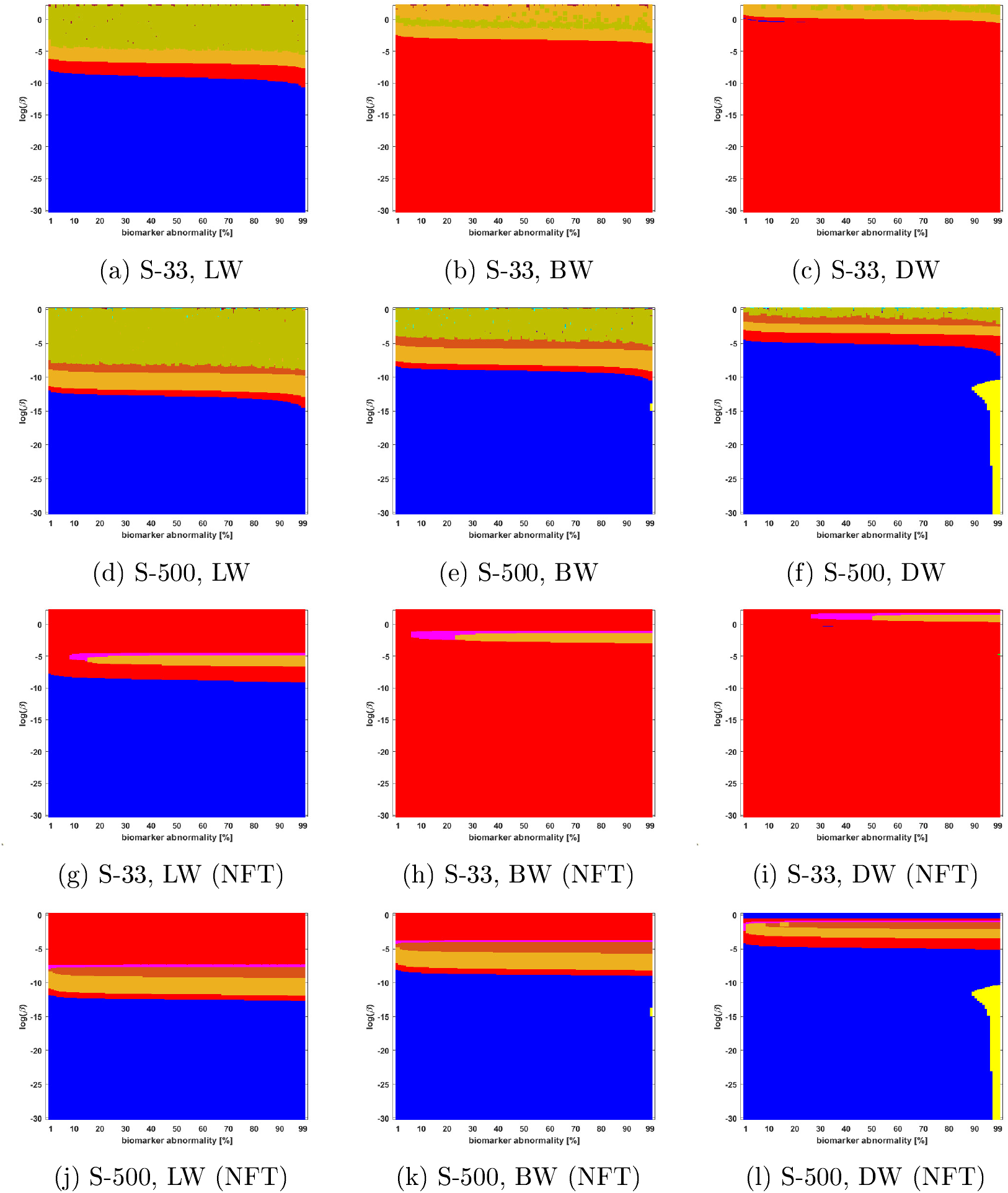
Observed computational (probabilistic) connectome *τ*P seed staging (top two rows) and *τ*P NFT staging (bottom two rows). Noise corrected backbone at a threshold of 1.28 with biomarker abnormality 1% ≤ *T* ≤ 100% (x-axis) and −30 ≤ ln(*β*) ≤ 0 (y-axis)

**Figure 6:**
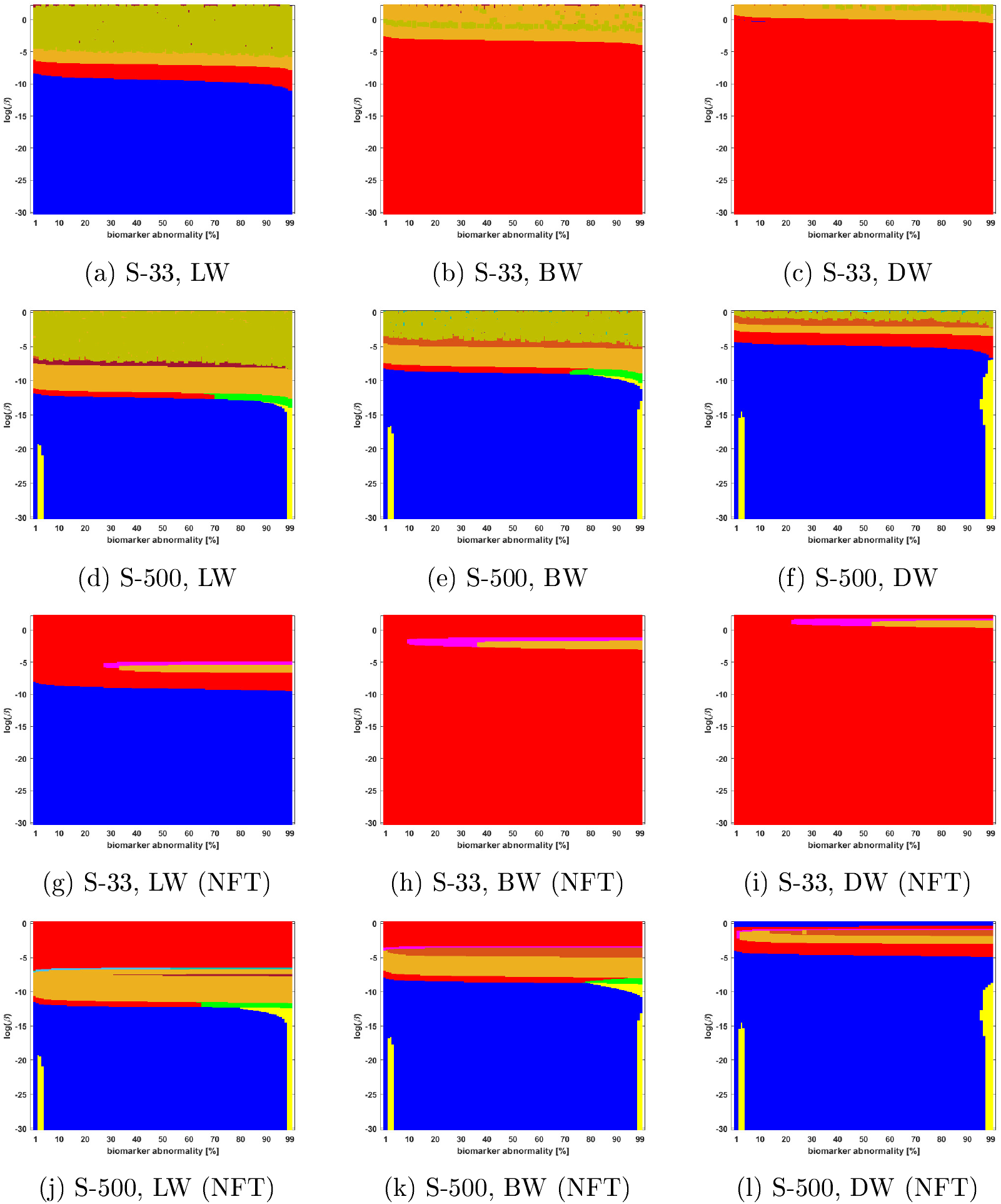
Observed computational (probabilistic) connectome *τ*P seed staging (top two rows) and *τ*P NFT staging (bottom two rows). Naive thresholding at a threshold of 5 × 10^−3^ with biomarker abnormality 1% ≤ *T* ≤ 100% (x-axis) and −30 ≤ ln(*β*) ≤ 0 (y-axis)

**Figure 7:**
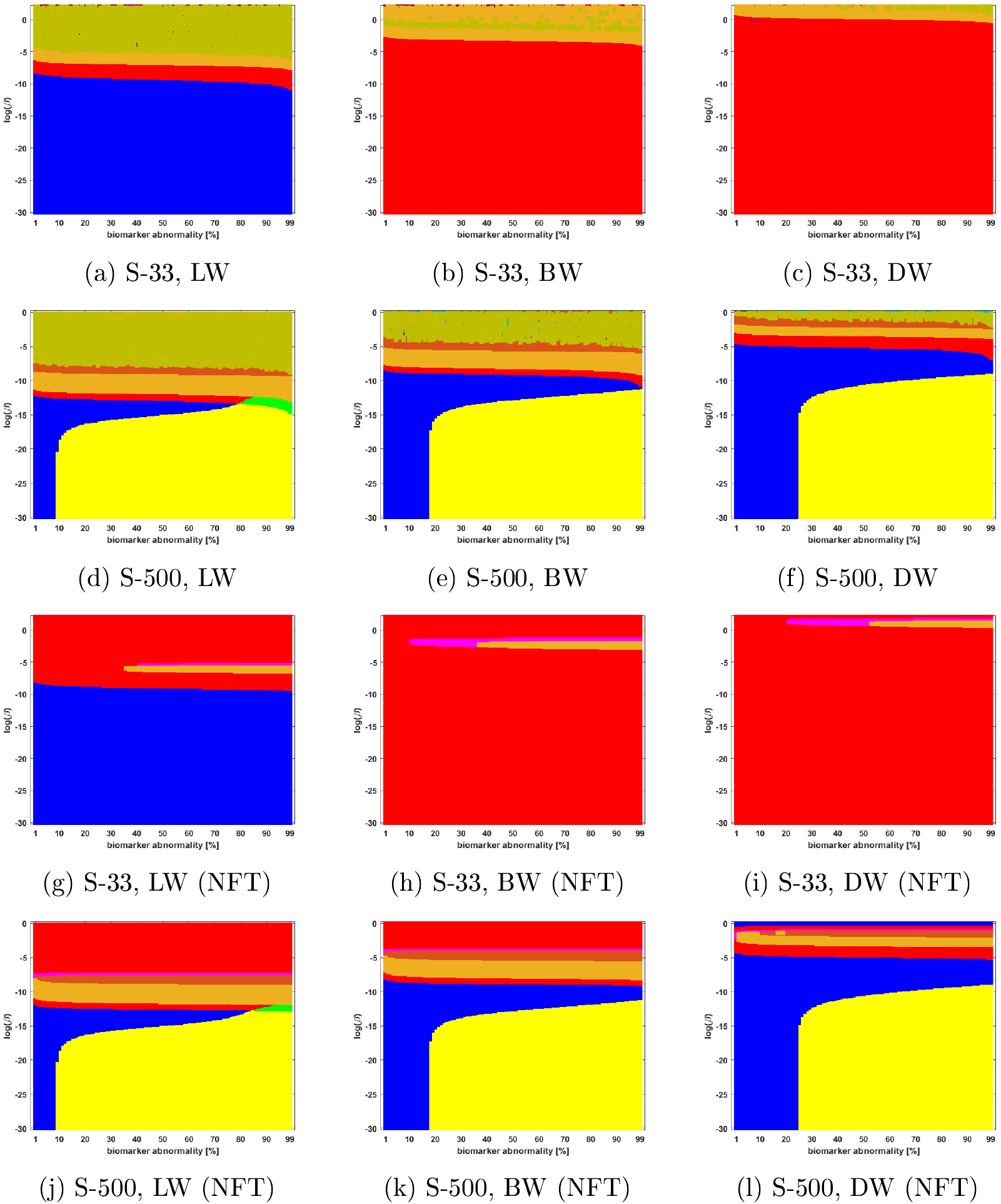
Observed computational (probabilistic) connectome *τ*P seed staging (top two rows) and *τ*P NFT staging (*δ* = 1, bottom two rows). Naive thresholding at a threshold of 1 × 10^−3^ with biomarker abnormality 1% ≤ *T* ≤ 100% (x-axis) and −30 ≤ ln(*β*) ≤ 0 (y-axis)

